# Parkinson’s disease risk factors are expressed at brain barriers

**DOI:** 10.1101/2025.10.20.683561

**Authors:** Floriane Bretheau, Océane Arevalo, Vincent Gélinas, Louise Reveret, Océane Martinez, Marlie Ali Yandza, Bastien Jacquet, Léa Ferreira, Morgan Bérard, Charles Joly Beauparlant, Steve Lacroix, Éric Boilard, Benoit Laurent, Frederic Calon, Martin Parent, Arnaud Droit, Aurélie de Rus Jacquet

## Abstract

Parkinson’s disease (PD) is characterized by the selective loss of dopaminergic neurons in the *substantia nigra pars compacta*, but whether its etiology is cell autonomous remains unclear. Increasing evidence implicates the blood-central nervous system (CNS) barriers in disease development, highlighting the importance of identifying genetic risk factors linked to cells forming the cerebrovasculature to advance this emerging area of research. The objective of this study is to identify PD genetic risk factors associated with blood-brain (BBB) and blood-cerebrospinal fluid (BCSFB) barriers, and to validate protein localization in human tissue and experimental models. To do so, we integrated genome-wide association studies and single nuclei RNA-sequencing datasets from the human postmortem substantia nigra (SN), midbrain, or cortical samples from control and PD donors. An in-depth bioinformatics analysis identified genes enriched in cell types that form the multicellular architecture of brain barriers, including *CAVIN2*, *ANXA1*, *ANO2*, and *LRP1B*. We further validated whether corresponding proteins were present in cell types associated with the blood-CNS barriers in human and mouse post-mortem tissues, as well as in iPSC-differentiated cells and choroid plexus organoids. Results showed that quantifying the proportion of endothelial cells expressing PD-related genes was under-evaluated at the transcript level compared to immunofluorescence analyses. In addition, we observed that CAVIN2 and ANXA1 proteins were more abundant at the vasculature of the substantia nigra vs. cortex, and CAVIN2 protein levels were reduced in PD vs. control human postmortem tissues. In contrast, the investigation of mouse postmortem samples demonstrated that the CAVIN2 protein is only present in a subset of mouse blood vessels, compared to nearly all vessels in human tissue. Similarly, mouse ANXA1 protein localizes to dopaminergic neurons of the substantia nigra and not at the vasculature, as seen in human tissue. The primary outcome of this study is the identification of PD-relevant risk genes specifically expressed at brain barriers and enriched in PD-relevant brain regions. The secondary outcome is the demonstration of poor transcript-protein correlation in – at least – a subset of PD risk factors, and a low interspecies conservation of protein localization for the selected candidates. In conclusion, the BBB and BCSFB may represent understudied contributors to PD, endothelial-specific proteins appear differentially regulated compared to transcripts, and experimental models require comprehensive validation to ensure relevance to the human condition.

---

Parkinson’s disease (PD) is the second most common neurodegenerative disorder after Alzheimer’s disease, affecting more than 11 million individuals worldwide in 2021, with a projected 50% increase in global incidence by 2035^1^. PD is a progressive disorder characterized by motor symptoms such as resting tremor, bradykinesia, and rigidity, along with non-motor manifestations including cognitive, psychiatric, and sleep disturbances^2^. Hallmark neuropathological features include the degeneration of dopaminergic neurons in the *substantia nigra pars compacta*^3,4^ and neuroinflammation^5–7^, and emerging reports propose that dysfunctional brain barriers may contribute to disease onset or progression^8–15^. Although its exact etiology remains unclear, PD is now recognized as resulting from the interaction of aging, genetic, and environmental factors^16^. Genetic factors account for approximately 5 to 10% of diagnosed cases^17^ and the largest meta-gene wide association study (GWAS) to date have identified more than 90 loci associated with increased PD risk, highlighting a wide range of genes and biological mechanisms involved in disease susceptibility^18,19^. Examples of risk factors include *STK39, GAK, SNCA, LRRK2, SYT11*, and *HIP1R*^20,21^, which are implicated in functional pathways across different cell types, such as neurons and glial cells^22^. In fact, the computational integration of GWAS and single cell or single nuclei RNA-sequencing datasets underscored the contribution of multiple cell types to PD risk, thus providing biological context to genetic risk factors. Oligodendrocytes and neurons are commonly identified in these studies, along with microglia and astrocytes which play key roles regulating cerebrovascular function^23,24^. However, the association of PD traits with other cell types that form brain barriers remains to be characterized.

The neuro-glia-vascular unit plays a key role in maintaining brain homeostasis through the integrity of the blood–brain barrier (BBB) and the blood–cerebrospinal fluid barrier (BCSFB)^25^. These barriers constitute essential physiological interfaces that maintain cerebral homeostasis. Indeed, they selectively regulate the exchanges of molecules and transmigration of immune cells between the blood and the brain parenchyma, and the BCSFB located at the choroid plexus (ChP) and meninges regulate exchanges between the blood and the cerebrospinal fluid (CSF). In PD, BBB alterations have been reported^26–29^ and recent clinical studies suggest a correlation between morphological changes at the ChP, motor symptom progression and dopaminergic degeneration^30,31^. The ChP produces CSF, and previous studies have documented a shift in the CSF molecular profile over the course of the disease^32,33^, further suggesting a possible implication of this tissue in disease progression. Therefore, brain barriers may contribute to PD pathophysiology and represent a potential target for the development of new therapeutic strategies^34^.

In this context, this study proposes an in-depth investigation of the PD-associated genes most relevant to cell types forming the BBB (e.g. endothelial cells, pericytes) and the BCSFB (e.g. epithelial and ependymal cells). We leveraged open access GWAS and single nuclei RNA-sequencing (snRNA-seq) datasets, and identified genes of interest including *ANXA1*, *ANO2*, *CAVIN2,* and genes of the low-density lipoprotein receptor family. We then confirmed protein expression and cellular localization in human postmortem brain and ChP tissue donated by people with PD or unaffected controls. Finally, to support future experimental and translational studies on these risk factors, their presence was validated in multiple relevant experimental models, including mouse and *in vitro* models derived from human induced pluripotent stem cells (iPSCs). Overall, we propose that PD-related genetic factors that affect brain barriers represent an underexplored opportunity to appreciate the complexity of PD etiology. However, it is cautioned that patterns of gene expression are not necessarily conserved across species, thereby warranting careful validation of experimental models to ensure future translation to patients.

## Material and methods

### Sex as a biological variable

Study design related to the human postmortem characterization of proteins of interest included male and female donors. However, PD is more prevalent in males vs females and the sample cohort reflects this sex bias. Study design related to the validation of proteins of interest in mouse and human iPSCs did not include sex as a biological variable, as previously published studies did not indicate sex bias in the presence/absence of the protein of interest in these models. These observations are corroborated by human postmortem data provided in this study. However, the biological function of these proteins may vary between male and female, and follow-up studies in experimental models may benefit from a sex-based stratification.

### Processing snRNA-sequencing study datasets

Bioinformatics analysis were performed on three publicly available datasets from Kamath et al., 2022 (GSE178265)^35^, Feleke et al., 2021 (GEO GSE178146)^36^ and Smajic et al., 2022 (GSE157783)^24^. These datasets captured the same cell types, including astrocytes, oligodendrocytes, pericytes, microglia, neurons, and endothelial cells. To study the BCSFB, we leveraged the dataset published by Yang et al ^37^ and extracted single nuclei gene expression data for endothelial, epithelial, mesenchymal, ependymal cells and monocytes. Each unique cell type was identified using established markers (**Supplementary Table 1**).

#### Data processing

The in-house RStudio package, SingleCell, which wraps and optimizes the main functionalities of the Seurat R package to analyze a Seurat object, was used to process the input data and form a clear analysis of the important markers per cell type. All matrices of RNA counts, features, and barcodes of the samples from the four different studies were extracted and placed into four distinct Seurat objects. For Yang et al., 2021, only the control samples for plexus choroid (*n = 6*) were used. For the other three datasets, all samples were used.

#### Data analysis

For each dataset, the in-house function SingleCell::analyze_integrated was performed based on the Seurat v4.4.0 package. First, a filtering step was applied, requiring a minimum of 100 gene features and 100 gene counts per features. To retain a maximum number of cells, no top threshold was used. Normalization was performed using Seurat::NormalizeData with the base parameters. Then, High Variable Genes (HVG) were identified using Seurat::FindVariableFeatures with the Variance Stabilizing Transformation (vst) argument with 2,000 features to stabilize the variance in cell expression. Furthermore, Seurat::ScaleData was used to scale the data with base parameters. 30 Principal Component Analysis (PCA) dimensions were used in the Seurat::RunPCA function to reduce the dimensionality and identify important patterns within the variation of cells expression. For each dataset, clusters were found with the base parameters of Seurat::FindClusters with a resolution of 0.2 to identify the base clusters for each dataset. Based on a set of key gene markers (**Supplementary Table 1**), clusters were assigned to a defined cell type, and PD-related genes of interest were mapped using a Uniform Manifold Approximation and Projection (UMAPs). Fold changes were calculated using the FindMarkers by Seurat with the following parameters: min.pct = 0.05, and logfc.threshold = 0.25.

#### Identification of arterial, capillary, and venous cells

To separate endothelial cells based on their zonation identity, clusters identified as endothelial cells were subdivided and analyzed using the base function of Seurat::FindSubCluster. Arterial cells were identified by the expression of *VEGFC* and *ALPL*, capillary cells by *MDSD2A* and *SCLA5*, and venous cells by *IL1R1* and *NR2F2*, and the threshold set at a minimum of log1p(0.5) expression.

### Association of PD genetic risk factors with cell types relevant to brain barriers

The gene list was extracted from the GWAS catalog v1.0.2.1 that compiled a total of 59 studies (https://www.ebi.ac.uk/gwas/docs/file-downloads)^38^. For each of the three snRNA-seq datasets, the GWAS gene list was reduced based on the genes found by the Average Expression base function from Seurat v.4.4.0 on the defined clusters^39^. For the Smajić et al. dataset, because the features were encoded in ENSEMBL, the GWAS gene list was translated with the base function mapIds from the AnnotationDb package version 1.64.1 with the database org.Hs.eg.db version 3.17.0 mentioning the ALIAS keytype^40^. To identify genes enriched in cell types of the neuro-glia-vascular unit and ChP under non-neurodegenerative conditions, the gene expression value for each cell type of interest was divided by the neuronal expression value for that particular gene. This analysis was carried out solely on datasets generated using control samples to avoid bias due to underlying neurodegenerative conditions. The top 100 genes enriched in endothelial cells were extracted and serve as the basis for comparing cell types of the neuro-glia-vascular unit. Genes with at least 10-fold higher expression level in cell types of interest vs. neurons were used to generate gene ontology enrichment data via Metascape^41^.

### Human postmortem brain tissues

Tissue samples were provided by the human brain bank at CERVO Brain Research Centre (Quebec, Canada), and written informed consent was obtained before donation (ethical approval provided by the Institutional Review Board CIUSSS de la Capitale-Nationale, project 2013-3 (146-2012, IUSMQ). The Université Laval Ethics Committee approved the brain collection procedures and the storing and handling of postmortem human brain material. Ethical approval for the use of human postmortem tissue was received from the Centre de Recherche du CHU de Quebec-Université Laval’s Institutional Review Board (project #2024-6837). Analyses were performed in accordance with the Code of Ethics of the World Medical Association (Declaration of Helsinki). Detailed demographics and clinical information on the postmortem brain samples is provided in Table 1. PD diagnosis was confirmed by a detailed neuropathologic examination that considered Lewy pathology consensus criteria^42–44^. These PD patients had no evidence of other cognitive, psychiatric or neurological disorders. In this study, postmortem human frontal cortex and substantia nigra (SN) tissue sections were examined for 5 control (1 female and 4 males, median age 81.4 years) and 5 PD donors (1 female and 5 males, median age 74 years). Postmortem delay (PMD) ranged from 3h to 23h for control donors and from 2.5h to 17h for PD donors. For human ChP samples, we examined sections from the lateral ChP of 4 control (2 females and 2 males, median age 73 years) and 4 PD donors (1 female and 3 males, median age 73 years). Postmortem delay (PMD) ranged from 3h to 18h for controls and from 15h to 36h for PD.

**Table 1.** Demographics and clinical data.

| Age, average (standard deviation) | Control : 81 (13.8) PD : 73,6 (6,6) |  |
| --- | --- | --- |
| Sex | Male: n = 5 (control), n = 7 (PD) Female: n = 2 (control), n = 2 (PD) |  |
| Disease status | Postmortem interval (h) | Cause of death |
| Control | 9 | Epilepsy |
| Control | 15 | Glioblastoma |
| Control | 23 | Cardiac insufficiency |
| Control | 3 | Medical assistance in dying |
| Control | 18 | Medical assistance in dying |
| Control | 10.5 | Stroke |
| Control | 3 | Septicemia |
| PD | 2.5 | Cardiac |
| PD | 17 | Septic shock |
| PD | 14 | Aspiration pneumonia |
| PD | 4 | Adenocarcinoma |
| PD | 12 | Aspiration pneumonia |
| PD | 36 | Aspiration pneumonia |
| PD | 15 | Cardiac |
| PD | 25 | Aspiration pneumonia |
| PD | 15 | Aspiration pneumonia |

### Culture of human iPSCs

The iPSC lines used in this study were reprogrammed from two different individuals. A first control line (female, line #1) was kindly provided by Prof. Dr. Thomas Gasser (Universitätsklinikum Tübingen) and Prof. Dr. Hans R. Schöler (Max-Planck Institute). This line served as the isogenic control of a LRRK2 G2019S pair, which was not investigated in this study. A second control line (female, line #2) was kindly provided by Prof. Dr. Tilo Kunath (University of Edinburgh). This line served as the isogenic control of an *SNCA* triplicated pair, which was not investigated in this study. These iPSC lines were cultured and maintained in growth medium consisting in 50% mTeSR Plus and 50 % TeSR E8 medium (StemCell Technologies, Vancouver, BC, Canada), passaged as small cellular aggregates using ReLeSR (StemCell Technologies), and plated onto Geltrex-coated culture dishes for maintenance (Thermo Fisher Scientific, Waltham, MA).

### Differentiation of iPSC-derived astrocytes

Control iPSCs lines were first differentiated into neural progenitor cells (NPCs) according to an established protocol^45^. Briefly, iPSCs were harvested with Accumax (Stemcell Technologies) and seeded at a density of 2×10^5^ cell/cm^2^ in 6 well plates pre-coated with Geltrex (FisherScientific, Hampton, NH) in mTeSR Plus medium supplemented with 10 µM Y-27632 (Selleck Chemicals). The iPSCs were then neuralized by dual SMAD inhibition to generate midbrain-patterned NPCs. Briefly, the cultures were sequentially exposed to LDN193189, SB431542, SHH(C25II), Purmorphamine and CHIR99021 until day 11 of the differentiation protocol, as described before^45^. On day 11, the cells were harvested using Accumax and seeded at a density of 7×10⁶ cells/well in 6-well plate pre-coated with Geltrex, in NPC medium (Neurobasal, 2% (v/v) B27, 1% (v/v) GlutaMAX, 20 ng/mL FGF2 (PeproTech, Cranbury, NJ), and 20 ng/mL EGF (Thermo Fisher) supplemented with 3 μM CHIR99021. On day 13, the medium was replaced daily with fresh NPC medium. On day 18, astrocyte differentiation was initiated by harvesting and seeding NPCs at 15,000 cells/cm^2^ in NPC medium supplemented with 10 µM Y-27632. The next day, the cells were washed with 1X PBS and maintained in astrocyte medium (ScienCell, Carlsbad, CA), followed by weekly passaging at a seeding density of 15,000 cells/cm² until day 28, as previously published^45^. This procedure yields 100% GFAP-positive astrocytes that have been extensively characterized in previous studies^46,47^.

### Differentiation of iPSC-derived ChP organoids and fixation

ChP organoids were generated using the STEMdiff Choroid Plexus Organoid Differentiation and Maturation kits (Stemcell Technologies, Vancouver, BC), as per the manufacturer’s instructions. Briefly, iPSCs were seeded at 9,000 cells/well in 96-well ultra-low attachment plates in seeding medium (organoid formation medium containing 10 µM Y-27632 (Selleckchem, Burlington, ON). Then, 100 µL of organoid formation medium was added on days 2 and 4. On day 5, the organoids were carefully transferred to 24-well ultra-low attachment plates containing 500 µL of induction medium. On day 7, the organoids were transferred on a parafilm sheet and embedded in 15 µL Matrigel (Corning, Glendale, AZ). After 30 minutes incubation at 37°C, Matrigel droplets containing organoids were transferred to a 6-well ultra-low adherent plate containing 3 mL of expansion medium. On days 10 and 13 medium was changed for choroid plexus differentiation medium, specific medium to induce ChP differentiation. On day 15, medium was changed for maturation medium and organoids were placed on an orbital shaker (50 RPM) located in a 37°C cell culture incubator. Medium was changed every other 2 to 3 days. After 52 days, the organoids were washed twice with phosphate buffer saline (PBS) and fixed overnight in 4% paraformaldehyde (PFA) at 4°C. Lastly, the organoids were washed twice with PBS and stored in PBS-0.02% sodium azide until cryosectionning.

### Animals

Male C57BL/6 and Slco1c1-icre/ERT2 mice were purchased from Charles River Laboratories or The Jackson Laboratory (JAX). All mice were housed in a facility that maintained temperature control at 23°C, a 12h/12h light-dark cycle, and *ad libitum* access to food and water. 3 to 5 months old mice were overdosed with a mixture of ketamine (400mg/kg) and xylazine (40 mg/kg) and transcardially perfused with PBS, and two subsequent procedures were employed to ensure optimal visualization of proteins of interest. To visualize Cavin2, mice were only perfused with PBS and brains were fresh frozen until tissue processing. To visualize ANXA1 and LRP1, PBS perfusion was followed by perfusion with 4% PFA, pH 7.4, in PBS. Brains were extracted, post-fixed for an additional 24h in 4% PFA at 4°C and then transferred into PBS+30% sucrose for at least 24h before tissue sectioning. Animal handling was performed in accordance with the guidelines of the Canadian Council on Animal Care and were approved by the Comité de Protection des Animaux du CRCHUQ-UL (protocols #CHU-2020-554; #CHU-2024-1574 and #CHU-24-1635).

### Isolation of mouse brain capillaries

The method used to generate microvessel-enriched extracts from fresh mouse brain samples has been described previously^48–51^. Briefly, this procedure consists in a series of centrifugation steps, including a density gradient centrifugation with dextran (Millipore sigma, Burlington, MA), after which the tissue is filtered through a 20-µm nylon filter. This generates two fractions: the material retained on the filter consists in cerebral microvessels (isolated microvessel-enriched fraction), whereas the filtrate consists in microvessel-depleted parenchymal cell populations. Isolated microvessels on the filter were resuspended in 3 ml of microvessel isolation buffer (HBSS; 15 mM HEPES, 147 mM NaCl, 4 mM KCl, 3 mM CaCl_2_, and 12 mM MgCl_2_) supplemented with 1% BSA and protease/phosphatase inhibitors, and spun at 2000 g for 10 min at 4 °C. The supernatant was discarded, and the pellet resuspended in 100 μl PBS. These cerebrovascular extracts were then deposited on glass slides (5 μl per slide) and left at room temperature (RT) for 30 min to allow adhesion. Afterwards, they were fixed using a 4% PFA solution in PBS, for 15 min at RT, and processed for immunostaining.

### ChP organoid and postmortem tissue processing

Organoids and mouse and human ChP tissues were embedded in Shandon™ M-1 Embedding Matrix or OCT (Thermo Fisher Scientific) with tissue sections cut at a thickness of 10 µm for organoids and mouse tissues, and at 18 µm for human tissues, using a cryostat (model CM3050S; Leica Biosystems, Concord, ON). All sections were collected directly onto Surgipath X-tra® slides which have a permanent positively charged surface (Leica Microsystems Canada) and stored at −20°C until use.

### Immunostaining

#### ChP organoids and postmortem tissue

ChP organoids, mouse and human tissues were labeled by immunofluorescence as follows. First, tissues were dried under vacuum at 4°C for 15 min. Human brain sections were exposed to white light for 4 days at 4 °C prior to tissue processing to reduce autofluorescence. Tissues were washed 3 times with KPBS (0.38% potassium phosphate dibasic anhydrous, 0.045% potassium phosphate monobasic anhydrous, 0.81 % NaCl in H_2_0 milliQ) for 8 min. For mouse and human tissues (except ChP), antigen retrieval was performed using a sodium citrate buffer at 85 °C for 15 min and cooled down for 30 min. Slices were then washed in KPBS (3 x 8 min) and incubated for 1 hour at RT in blocking buffer (5% donkey serum (Millipore sigma), 0.1% Triton-X (Millipore Sigma) in KPBS). Tissues were washed three times with KPBS for 8 minutes each, and then incubated overnight in the primary antibody solution at 4°C. Primary antibodies used in this study are listed in Supplementary table 2 and were used at the following dilutions: rabbit anti-Annexin 1 (1:100, Invitrogen, 71-3400), rabbit anti-Annexin 1 (1:200, Abcam, ab214486), rabbit anti-ANO2 (1:200, Abcam, 113443), mouse anti-AQP1 (1:100, Santa Cruz Biotechnology, SC-32737), goat anti-collagen IV (1:500, Millipore Sigma, AB769), mouse anti-GFAP (1:200, Novus Biologicals, NBP1-05197), rabbit anti-SDPR (CAVIN2) (1:200, Bio-Techne, NBP2-57218), rabbit anti-SDPR (CAVIN2) (1:200, Proteintech, 12339-1-AP), rabbit anti-LRP1 (1:200, Abcam, Ab92544). The next day, the slices were washed in KPBS (3 x 8 min) and incubated for 2.5 hours in Alexa-conjugated secondary antibodies (1:250 to 1:500 dilution, in blocking buffer). After incubation, the slices were washed three times in KPBS, incubated for 10 min in a DAPI nuclear stain solution (1:5,000, in KBPS) and washed again three times in KPBS. Human tissues were incubated in 0.1% Sudan Black solution prepared in 70% ethanol (Greenfield Global, Toronto, ON). Finally, slices were mounted using Fluoromount-G medium (Fisher Scientific, Waltham, MA).

#### iPSCs-derived astrocytes

iPSC-derived astrocytes were seeded in 48-well plates (40,000 cells/cm²) on coverglass pre-coated with Geltrex (Fisher Scientific). After 24 hours of incubation, cells were washed twice with Dulbecco’s PBS (DPBS) and fixed for 20 minutes in a 4% PFA solution under a chemical hood. Cells were then washed twice with DPBS and incubated for 1 hour at RT in a blocking/permeabilization solution (10% (v/v) FBS, 1% (w/v) BSA and 0.3% (v/v) Triton-X in DPBS). After two additional DPBS washes, the cells were incubated overnight at 4 °C in the primary antibody solution consisting of 1% (w/v) BSA in DPBS. The following antibodies were used: rabbit anti-Annexin 1 (1:200, Invitrogen, 71-3400), rabbit anti-Annexin 1 (1:200, Abcam, ab214486), rabbit anti-SDPR (CAVIN2) (1:200, Bio-Techne, NBP2-57218), rabbit anti-SDPR (CAVIN2) (1:200, Proteintech, 12339-1-AP), rabbit anti-ANO2 (1:200, Bio-Techne, NBP2-92888), rabbit anti-ANO2 (1:200, Abcam, 113443), mouse anti-GFAP (1:500, Novus Biologicals, NBP1-05197). The next day, cells were washed twice with DPBS and incubated for 1 hour at RT in Alexa-conjugated secondary antibodies (1:1,000 dilution, 1% BSA in PBS). Lastly, cells were washed twice in DPBS and incubated for 5 minutes at RT in a DAPI nuclear stain solution (1:5,000 dilution). Coverglasses were then mounted onto slides with a drop of ProLong™ Diamond Antifade Mountant. Slides were dried at RT protected from light for 24 hours. Coverslip edges were sealed with clear nail polish, and slides were stored at 4 °C until image acquisition.

#### Mouse brain capillaries

All samples were blocked and permeabilized for 1 h with PBS containing 10% (v/v) Normal Horse Serum (NHS) and 0.2% (v/v) Triton X-100. Incubation with primary antibodies (rabbit anti-SDPR (CAVIN2) (Proteintech, 12339-1-AP), goat anti-collagen IV (Merck Millipore, AB769) was performed overnight at 4 °C in PBS with 1% (v/v) NHS and 0.05% (v/v) Triton X-100. After three washes, the samples were incubated for 1 hour at RT in Alexa-conjugated secondary antibodies (1:1,000 dilution, 1% Normal Horse Serum in PBS). Slides were then incubated in a DAPI nuclear stain solution for 10 min at RT, and mounted with mounting medium (VectaMount AQ Mounting Medium).

### Confocal microscopy

Tissue sections and astrocyte monolayers were imaged on a Zeiss LSM 800 confocal microscope system equipped with 405-nm, 499-nm, 561-nm and 640-nm lasers. Confocal images were acquired using the Zen Blue Edition software (version Zen Blue 3.9, Zeiss, Oberkochen, Germany).

### Image analysis

The quantification of vascular-specific CAVIN2 signal intensity was performed using Fiji (v. 2.9.0). First a mask was created using the collagen IV channel by (i) applying a threshold (Huang method) and (ii) removing background noise using the “particle analysis” and “remove outliers” (radius=2) tools. Next, the regions of interest (ROI) created from the collagen IV mask were overlayed onto the CAVIN2 channel, and CAVIN2 signal intensity located within the ROIs was measured. For large vascular segments (e.g. arterioles), ROI selection was manually curated to ensure best fit. To ensure that the final quantifications reflect vascular signal, particles smaller than 5 µm^2^ were excluded from analysis.

### Statistical analysis

Data was analyzed using GraphPad Prism version 10 (La Jolla, CA). Data encompassing multiple groups were analyzed using one-way or two-way ANOVA with multiple comparison correction (**Fig. 1, 3, 4**).

**Figure 1.**
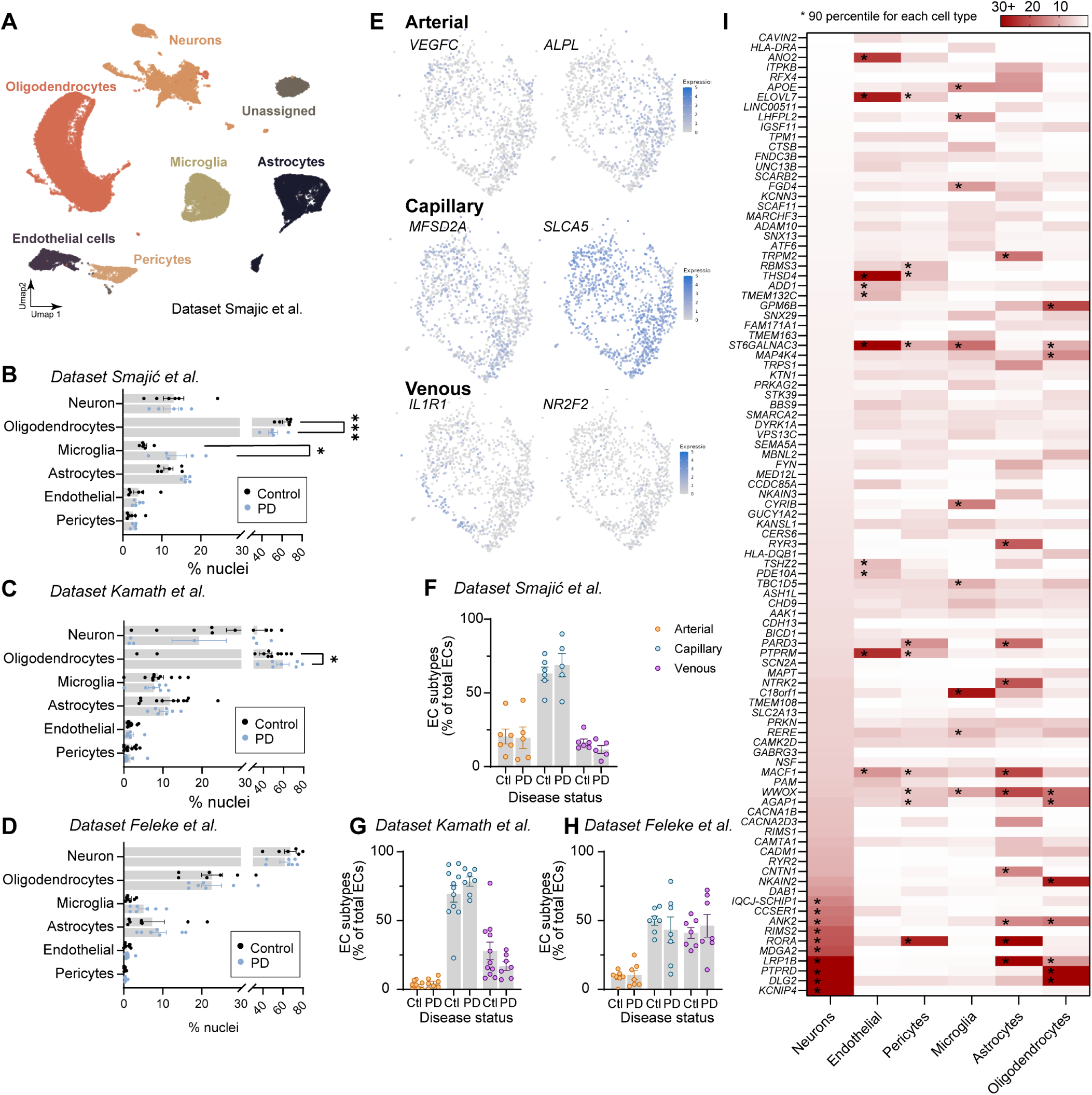
snRNA-seq datasets captured cell types of the neuro-glia-vascular unit. **A**. UMAP illustrating cell type clusters for the dataset by Smajic et al. **B-D.** Graphs show the number of nuclei for each cell type included in the study and obtained from Smajić et al. (B), Kamath et al. (C), and Feleke et al. (D). **E**. UMAP showing endothelial expression of gene markers of arterial, capillary and venous cells (Smajić et al. dataset). **F-H.** Graphs represent the proportion of arterial, capillary, and venous endothelial cell nuclei in control and PD samples, for Smajić et al. (F), Kamath et al. (G), and Feleke et al. (H). **I.** Heatmap shows gene expression for PD risk factors (Smajić et al. dataset) for each cell type. Genes are organized from the least to the highest expressed in neurons. * Genes in the 90 percentile for each cell type. Abbreviation: Ctl, control; ECs, endothelial cells; PD, Parkinson’s disease.

### Data availability

SnRNA-seq datasets used in this study were published in publicly available repositories, and they were obtained using accession numbers GSE178265, GSE178146, and GSE157783. Custom code generated to analyze the snRNA-seq data will be made available upon publication.

## Results

### PD risk factors are enriched in cell types forming the neuro-glia-vascular unit

To identify PD risk factors enriched in cells forming the neuro-glia-vascular unit, we leveraged three publicly available snRNA-seq datasets generated from human postmortem tissue of the *substantia nigra* (SN), midbrain, and anterior cingulate cortex of controls and PD donors. The cell types of interest were identified by marker genes previously established and include endothelial cells, pericytes, microglia, and astrocytes (**Supplementary Table 1**). In addition, oligodendrocytes were included in the analysis to serve as a reference glial cell type not associated with the neuro-glia-vascular unit. Neurons were included to enable the identification of PD risk factors that are not only enriched in cell types associated with brain barriers, but also display low expression in neurons. In the perspective of producing a meta-analysis, we attempted to integrate these three datasets as a single Seurat object and correct for potential batch effects that would confound data analysis. Integration was performed with an in-house package based on Seurat workflow that normalizes and scales data before finding common anchors (gene modules) between all three datasets. However, due to substantial batch effect, the datasets could not be integrated and were consequently analyzed separately. The snRNA-seq dataset published by Smajić et al. originates from midbrain samples collected from 6 control and 5 idiopathic PD donors. The Kamath et al. dataset includes the SN from 8 control and 10 PD donors, while the Feleke et al. dataset comprises anterior cingulate cortex from 7 control and 7 PD donors. Data UMAP separated cell clusters and confirmed that datasets captured cell types of the neurovascular unit (**Fig. 1A**). Here, we focused our study on cell types involved in BBB maintenance and regulation (i.e., endothelial cells, pericytes, astrocytes, and microglia), as well as oligodendrocytes and neurons. Nuclei quantification showed that oligodendrocytes represent the most abundant cell population in datasets originating from the midbrain and SN (**Fig. 1B-C)**, while neurons are the most abundant nuclei in the dataset produced using cortical tissue (**Fig. 1D**). Nonetheless, endothelial cells are represented in all three studies, and they can be further separated by zonation along the vascular arborescence to identify arterial (*VEGFC*, *ALPL*), capillary (*MFSD2A*, *SLCA5*) and venous (*IL1R1*, *NR2F2*) endothelial cells (**Fig. 1E**). Data showed that capillary endothelial nuclei are over-represented in the midbrain and SN samples, while nuclei of venous identity are more abundant in the cortical sample (**Fig. 1F-H**).

To identify PD risk factors relevant to each cell type of the neuro-glia-vascular unit, the expression level of GWAS-relevant genes was calculated for endothelial cells, pericytes, microglia, astrocytes, oligodendrocytes, and neurons originating from control donors (**Fig 1I**). We reasoned that the most relevant genes would be those with the highest fold expression when compared to neuronal expression, and we restricted this analysis to control samples to avoid biases and misrepresentation of cell populations lost or increased over the course of PD. This strategy led to the identification of Caveolae Associated Protein 2 (CAVIN2), Anoctamin 2 (ANO2) and Annexin A1 (ANXA1) as the top 3 enriched genes in endothelial cells vs. neurons. Other enriched genes include *GJA5* (pericytes), the Major Histocompatibility Complex (MHC) Class II family (*HLA-DRB5*, *HLA-DRB1*, *HLA-DRA*; microglia), *ITPKB* (astrocytes), *CD38* (pericytes, astrocytes), and *DMRT2* (oligodendrocytes) (**Fig. 2A-E**). Of note, the pattern of GWAS-related gene expression appears closely related between oligodendrocytes and neurons, and between endothelial cells and pericytes. The analysis of gene ontology (GO)-relevant pathways specific to endothelial cells and pericytes revealed that PD risk factors associated with these cell types (i.e. 10-fold enriched vs. neuronal expression) are implicated in immune responses, transport, cell activation, and vascular function (**Fig. 2F**).

**Figure 2.**
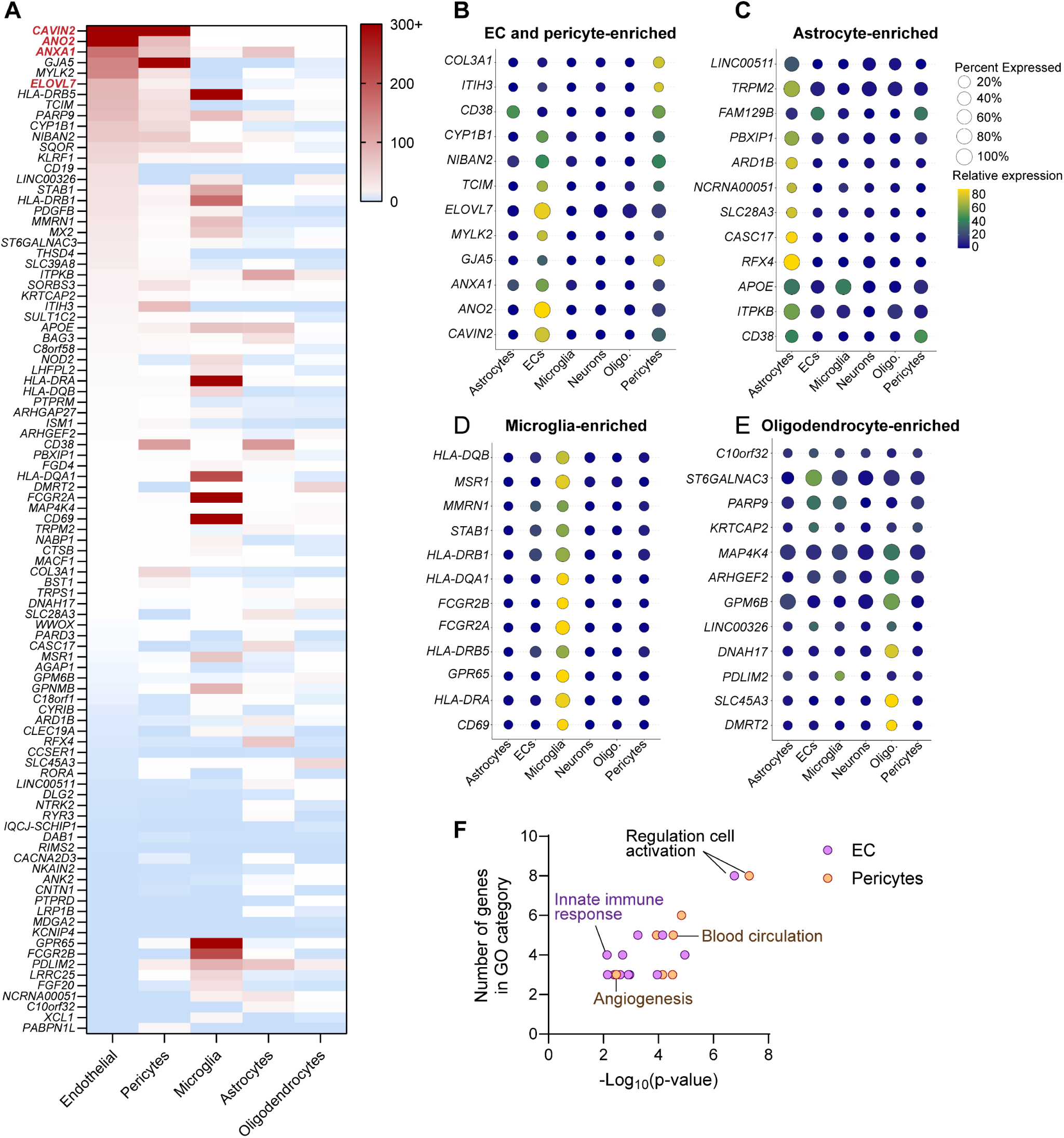
Integration of GWAS and snRNA-seq reveals PD risk factors genes enriched at the neuro-glia-vascular unit. **A**. Heatmap shows PD risk factors enriched in each cell type (Smajić et al. dataset). The scale is fold-change expression relative to neurons. Gene names in red indicate PD risk factors further investigated in this study. **B-E.** Dot plots represent the percent and relative expression of PD risk factor enriched in endothelial cells (ECs) and pericytes (B), astrocytes (C), microglia (D), and oligodendrocytes (E). **F**. Gene ontology analysis reveals pathways most affected by PD risk factors enriched in ECs and pericytes.

### Altered expression of the caveolae-associated CAVIN2 protein at the PD brain vasculature

Among the PD risk factors enriched at the neuro-glia-vascular unit, *CAVIN2*, *ANO2* and *ANXA1*, are the most expressed in brain endothelial cells. To confirm expression and localization of the encoded proteins, human postmortem tissue from control (n= 5) or PD (n= 5) donors was sectioned from the frontal cortex and SN, and immunostained for fluorescence microscopy. The vascular enrichment of CAVIN2 is demonstrated by the colocalization with Collagen IV in both the cortex and SN of control and PD patients (**Fig. 3A, B, Supplementary Fig. 1A-B**). Furthermore, a quantitative analysis revealed a significant decrease in CAVIN2 intensity in the cortex and SN of PD donors compared to controls (****p < 0.0001) (**Fig. 3C**), suggesting disease-related alterations at the protein level that were not observed in snRNA-seq studies. The quantification also revealed that CAVIN2 is enriched in the SN vs. cortex, further supporting the relevance of this risk factor to the regional vulnerability observed in PD pathology. Signal specificity was confirmed using a second antibody **(Supplementary Fig. 1C**). When comparing the pattern of protein vs. gene expression, data revealed that up to 96% of brain vessels were positive for CAVIN2 by immunostaining, but only 38 to 78% endothelial nuclei expressed *CAVIN2*, with variability depending on the snRNA-seq dataset (**Fig. 3D, E**). Lastly, we confirmed that expression of this PD-relevant gene is conserved across species, albeit at lower levels in mouse vs. human tissue. Immunofluorescence images spanning the wild-type mouse cortex and SN, as well as isolated mouse capillaries, confirmed vascular CAVIN2 signal in some (but not all) vascular segments (**Fig. 3F, G, Supplementary Fig. 1D, E**).

**Figure 3.**
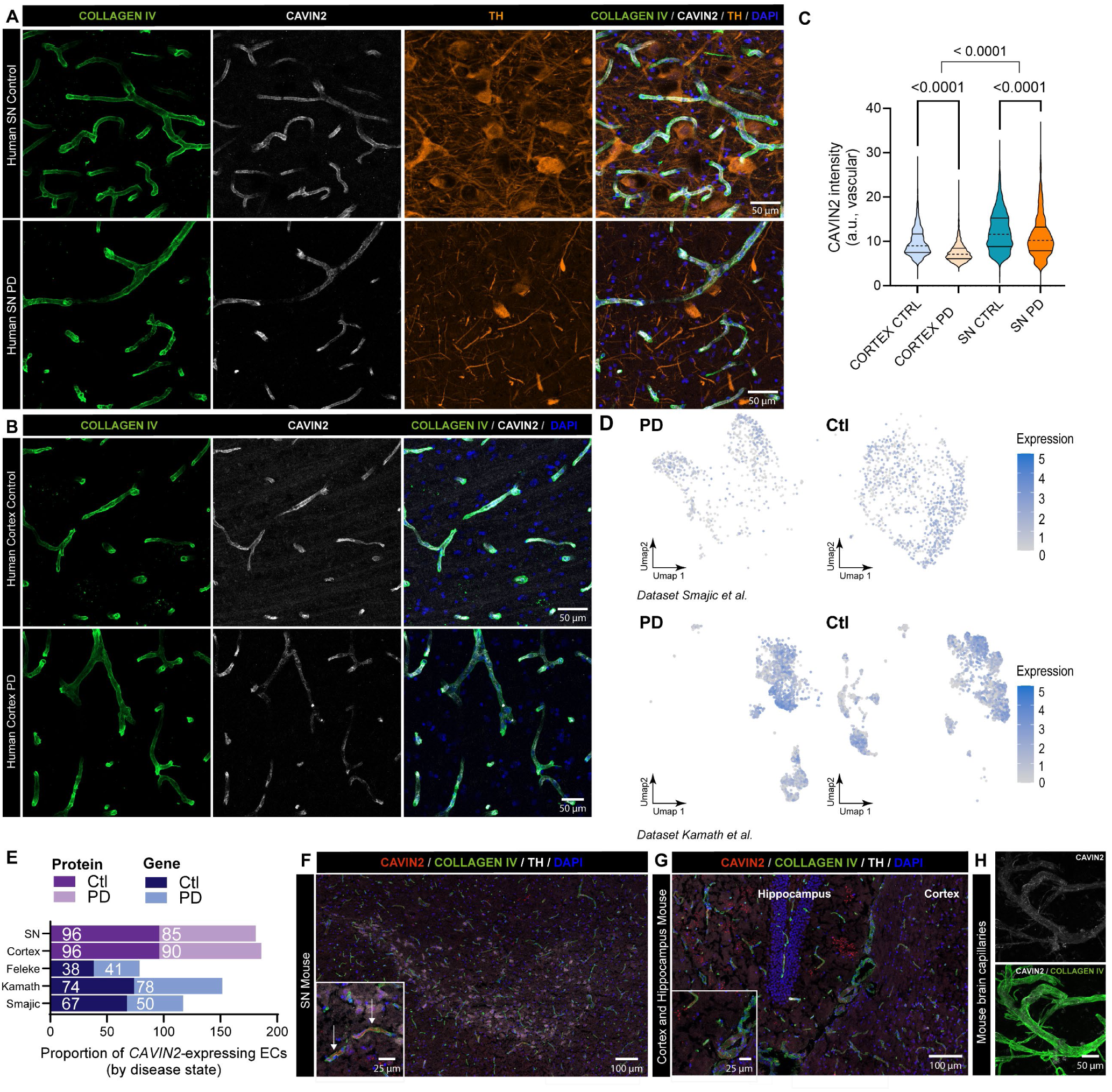
CAVIN2 is expressed by brain endothelial cells and protein level is altered in PD brains. **A-B.** Representative confocal images showing CAVIN2 (white), Collagen IV (green), TH (orange) expression in the SN (A) and frontal cortex (B) of human postmortem samples from controls and PD donors. **C.** Quantification of CAVIN2 signal intensity in collagen IV-positive blood vessels associated with acquisitions A and B. The violin plot shows the median (dashed line) and interquartile range (solid line). Data are from five patients and five control donors (n˃ 1,510 vessels analyzed for each condition). Statistical analysis was performed using two-way ANOVA with Tukey’s multiple comparisons correction. **D.** UMAPs represent *CAVIN2* expression level in endothelial nuclei related to Smajić et al. and Kamath et al. **E.** Graph representing the proportion of *CAVIN2*-expressing endothelial nuclei by snRNA-seq (blue bars), and CAVIN2-positive vascular fragments by immunofluorescence (purple bars). **F-G.** Representative confocal images showing the presence of CAVIN2 (red), Collagen IV (green), TH (white) in blood vessels of the mouse SN **(F)**, cortex and hippocampus **(G**). White arrows point to CAVIN2-positive blood vessels. (**H**) Representative confocal images showing the presence of CAVIN2 (white) and COLLAGEN IV in mouse brain capillaries. Abbreviations: Ctl, control; PD, Parkinson’s disease.

### The cell type-specificity of ANXA1 in the cortex and SN is not conserved across species

The investigation of ANXA1 in human postmortem cortical and SN tissue by immunofluorescence confirmed a strong vascular localization, along with a presence in discrete astrocyte populations (**Fig. 4, A-B, H, Supplementary Fig. 2A-B**). The quantification of ANXA1 at the vasculature did not reveal differences between control and PD donors, however fluorescent signal was higher in the SN compared to the cortex (**Fig. 4C)**. However, the pattern of gene vs. protein expression showed that the proportion of *ANXA1*-expressing endothelial nuclei is lower than protein abundance *in situ* (**Fig. 4D-E**). To support future investigations and confirm that this PD-relevant protein has a similar cellular localization across species, we obtained wild-type mouse brain samples and performed immunofluorescence staining for ANXA1, using the same antibody. Images show that mouse ANXA1 is enriched in dopaminergic neurons of the SN, and the protein was not reliably detected in the cortex or hippocampus (**Fig. 4F-G, Supplementary Fig. 2C-D**), suggesting that the pattern of ANXA1 protein localization is not conserved across species. Furthermore, we attempted to characterize ANO2 as a potential risk factor of interest and performed immunostaining of human postmortem tissue. The resulting images revealed the presence of occasional astrocytic staining in the cortex, but there are few resources available to investigate this protein and signal specificity could not be validated by the second selected antibody **(Supplementary Fig. 3A**). Furthermore, the expression and cellular localization of ANO2 could not be validated in mouse brain sections, as available antibodies yielded poor image quality. Nonetheless, we leveraged translational human cell culture models to validate the presence of proteins of interest in these platforms. The ANXA1 and ANO2 proteins were selected for this analysis, as they were identified in discrete astrocyte populations of the human postmortem brain. Primary astrocytes as well as two independent control iPSC-derived astrocyte lines were immunostained, and the images confirm ANXA1 and ANO2 expression in these cultures using two different antibodies **(Fig 4I-J; Supplementary Fig. 2E-F; Supplementary Fig. 3**). Lastly, ANXA1 signal specificity was confirmed using a second primary antibody **(Supplementary Fig. 2G**). Overall, these results support the future investigation of CAVIN2 and ANXA1 as neurovascular PD risk factors potentially implicated in disease neuropathology, and species differences in protein localization should be considered when selecting experimental models. The development of new resources would be valuable efforts to further explore the contributions of ANO2 to PD pathology.

**Figure 4.**
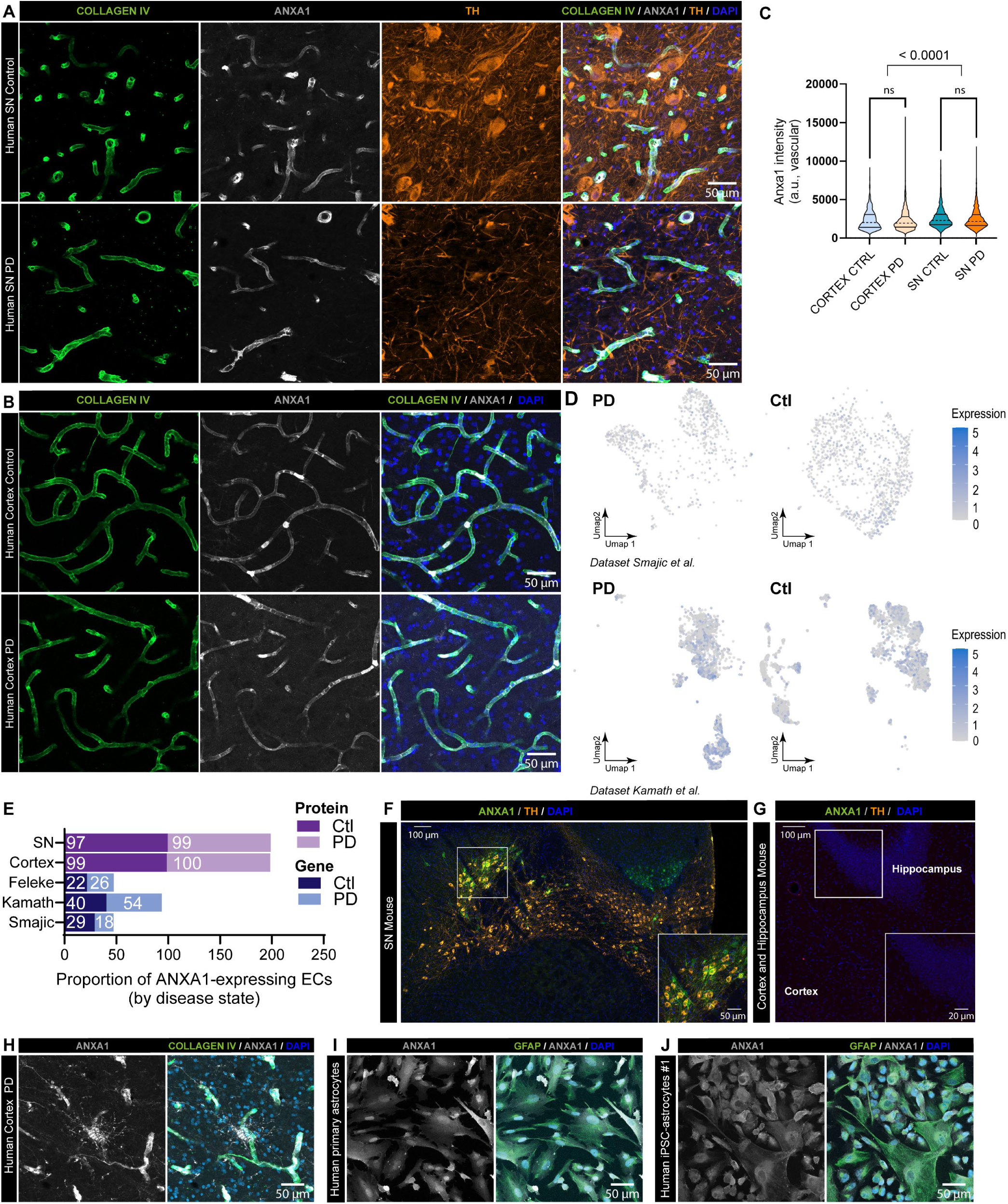
ANXA1 is localized in human blood vessels and astrocytes, and in mouse dopaminergic neurons. **A-B.** Representative confocal images showing ANXA1 (white), Collagen IV (green), TH (orange) levels in blood vessels of the SN (A) and cortex (B) in controls and patients with PD. **C.** Quantification of ANXA1 signal intensity in blood vessels associated with acquisitions A and B. The violin plot shows the median (dashed line) and interquartile range (solid line). Data are from five patients and five control donors (n ˃ 1,070 vessels analyzed for each condition). Statistical analysis was performed using two-way ANOVA with Tukey’s multiple comparisons correction (ns: non significant). **D.** UMAPs represent *ANXA1* gene expression level in endothelial nuclei. **E.** Graph represents the proportion of *ANXA1*-expressing endothelial nuclei by snRNA-seq (blue bars), and ANXA1-positive vascular fragments by immunofluorescence (purple bars). **F-G.** Representative confocal images showing ANXA1 (green) and TH (orange) in the mouse SN (F) and in the cortex and hippocampus (G). **H-J.** Representative confocal images showing ANXA1 (white) and Collagen IV or GFAP (green) in postmortem human cortical astrocytes (H), primary human astrocytes (I), and iPSC-derived astrocytes (J). Abbreviations: Ctl, control; ECs, endothelial cells; PD, Parkinson’s disease; SN, substantia nigra.

### PD risk factors are enriched in cell types forming the BCSFB at the ChP

Despite evidence demonstrating that aging and neurodegenerative diseases induce both morphological and functional changes at the ChP ^31,52–55^, this brain structure has been largely overlooked in PD-focused studies. Therefore, we mirrored our bioinformatics analysis of the neuro-glia-vascular unit using a published snRN-seq dataset of the human ChP^56^ (**Fig. 5A-B**, **Supplementary Table 1**). We identified PD risk factors most enriched in endothelial, ependymal, epithelial, mesenchymal cells and macrophages that could inform disease etiology (**Fig. 5C**). Among these, *ANO2*, *ELOVL7*, and *ANXA1* are genes enriched at both the neuro-glia-vascular unit and the ChP, therefore they were selected for in-depth validation. In addition, *LRP1B* was identified as a risk factor of interest. It is a member of the low-density lipoprotein (LDL) receptor family, which consists of 7 structurally related proteins, among which features the LRP1 receptor implicated in α-synuclein transport and propagation^57^. To validate the presence of these gene products at the ChP and ependymal regions of the brain, we prepared samples originating from the lateral ventricle of human and mouse postmortem tissues, and generated ChP organoids derived from control iPSCs (**Fig. 5D-E**). First, the morphological integrity of each ChP tissue was validated by immunostaining using established ChP markers (**Fig. 5F-H)**. Then, we assessed the anatomical localization of the proteins of interest. Among the candidates, ELOVL7 and LRP1B were not further pursued because the commercially available antibodies tested in this study yielded poor image quality. However, high quality and validated antibodies are commercially available to study LRP1, which is structurally related to LRP1B and a protein of interest for its role in α-synuclein propagation. Therefore, LRP1 immunostaining was performed on human postmortem tissue from control (n= 4) or PD (n= 4) donors **(Fig. 6A, Supplementary Fig. 4A-B)**, and the images show protein expression in the epithelial and stromal layers of the ChP. The presence of LRP1 at the mouse ChP was also confirmed **(Fig. 6B, Supplementary Fig. 4C)** and immunostaining of human iPSC-derived ChP organoids validated that this advanced *in vitro* model can be leveraged to study this protein **(Fig. 6C)**.

**Figure 5.**
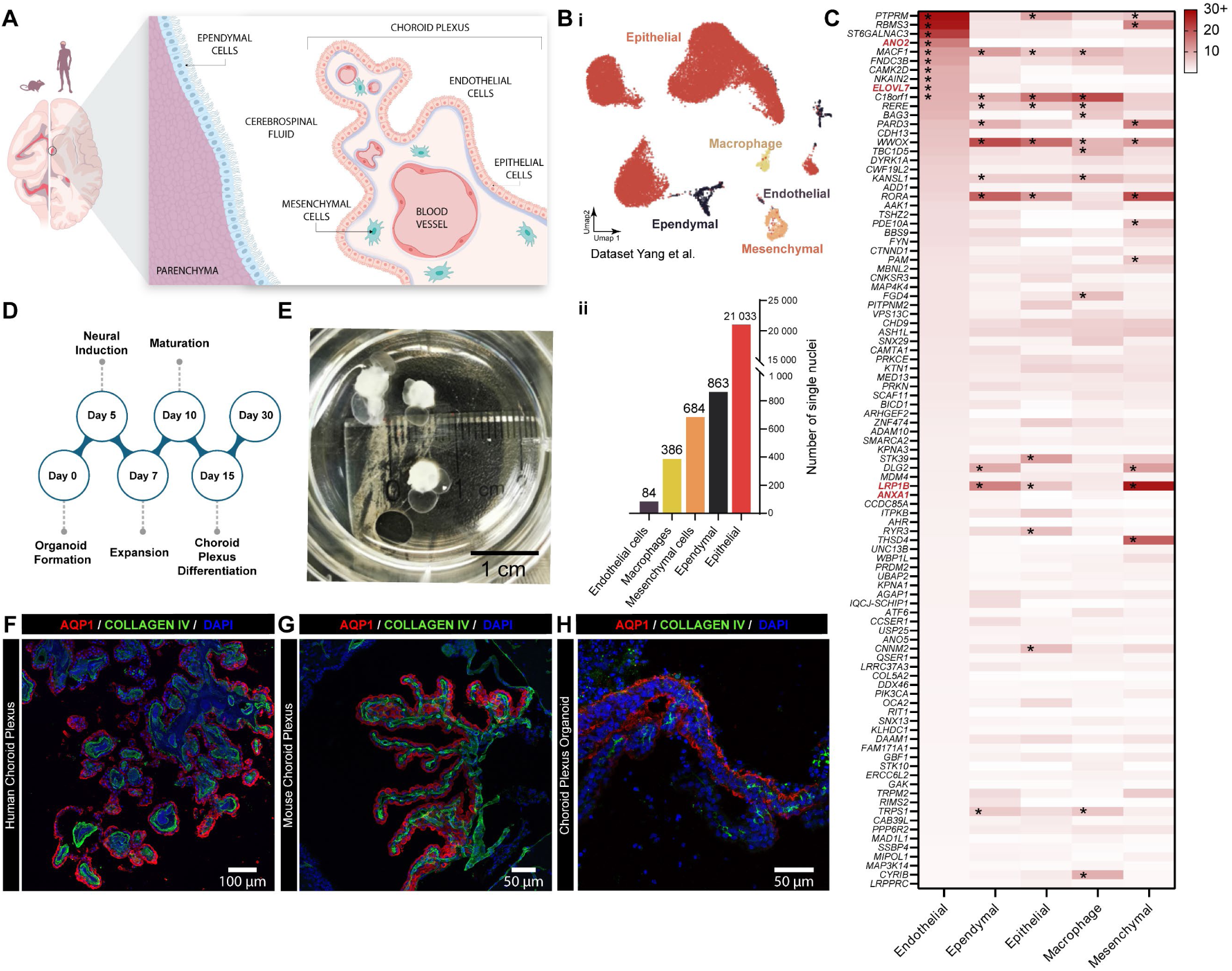
Identification of PD risk factors expressed at the BCSFB. **A.** Schematic localisation and cellular organisation of the ChP in mouse and human brains. **B. i.** UMAP represents snRNA-seq clusters that separate ChP cell types. **ii.** Graph shows the number of nuclei identified in the snRNA-seq study by Yang et al. **C.** Heatmap shows cell type-specific gene expression of PD risk factors. * Gene in the 90 percentile for each cell type. Gene names in red indicate PD risk factors investigated in this study. **D.** Timeline of ChP organoid differentiation from human iPSCs. **E.** Representative image of a mature CSF-producing ChP organoid fixed in 4% PFA at day 30 of differentiation. **F-H.** Representative confocal images showing APQ1 (red) and Collagen IV (green) in human (**F**), mouse (**G**) and organoid (**H**) ChP.

**Figure 6.**
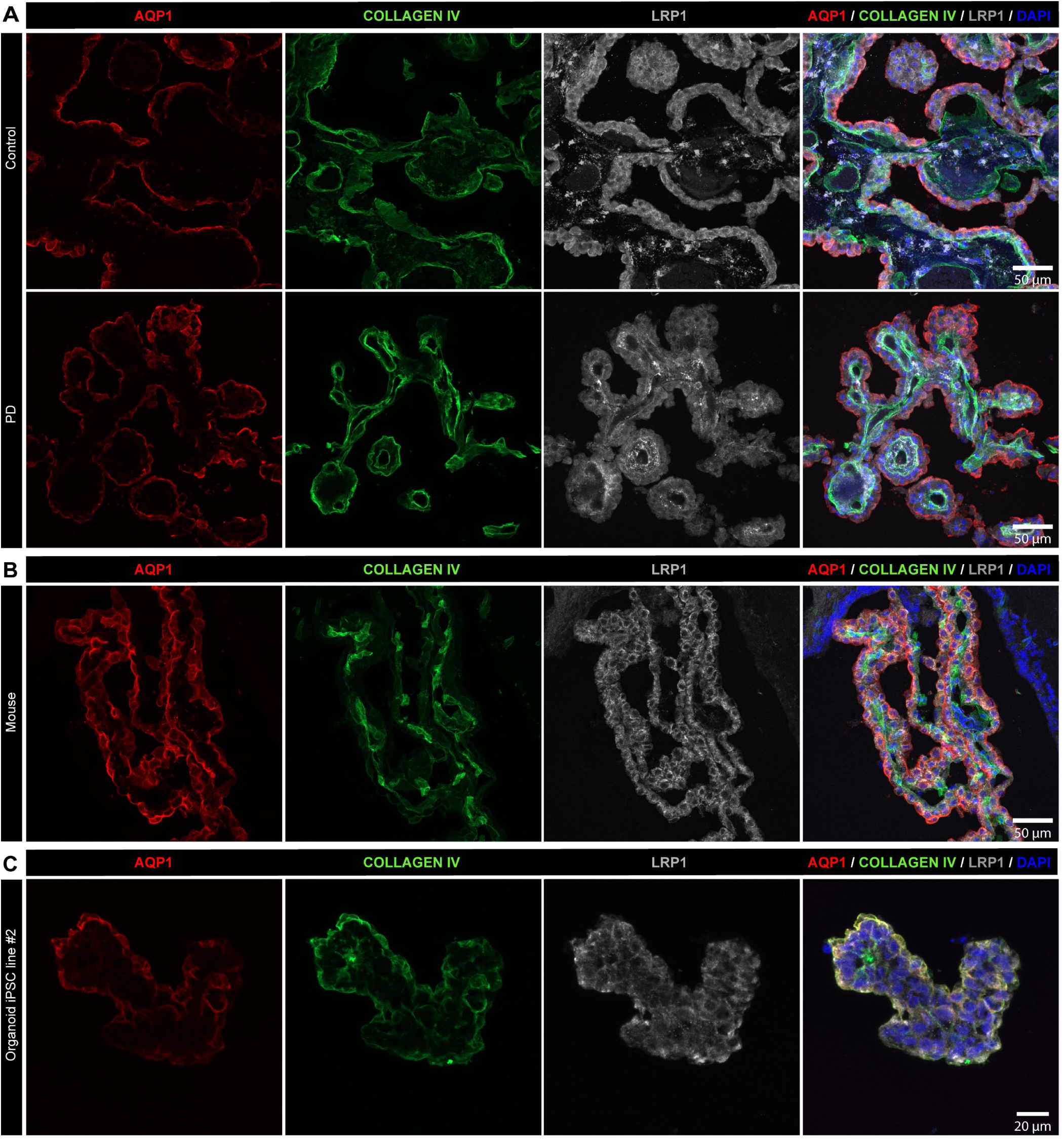
LRP1 is present at the ChP and cellular localization is conserved across species. **A-C.** Representative confocal images showing LRP1 (white), AQP1 (red) and Collagen IV (green) in ChP tissue from human control and PD postmortem tissues (**A**), mouse (**B**) and ChP organoid (**C**).

### The ependymal layer is enriched in ANXA1 and ependymal expression is conserved across species

We visualized the ANXA1 protein by immunostaining human postmortem tissue of the lateral ventricle/ChP from control and PD donors, and observed the presence of this protein at the ependymal layer (**Fig. 7A**). This cellular lining of the brain ventricles forms a protective barrier with the brain parenchyma, and ANXA1 immunostaining was observed in both control (n= 4) and PD (n= 4) human postmortem tissues (**Supplementary Fig. 5A-B**). Then, we confirmed that experimental models recapitulate human-based observations. Notably, the mouse ependymal layer is also enriched in ANXA1 (**Fig. 7B; Supplementary Fig. 5C**), and the protein was observed within the cellular soma and neuroepithelial cilia. However, *ANXA1* gene expression in epithelial nuclei did not mirror protein signal (**Supplementary Fig. 5D**). Furthermore, ChP organoids recapitulated a cellular structure that highly resembled the ANXA1 expressing ependyma (**Fig. 7C**). Signal specificity was confirmed using a second antibody across all sample types **(Supplementary Fig. 6**). In addition to LRP1 and ANXA1, ANO2 was also explored as a PD-relevant protein potentially expressed at the ChP. The ANO2 antibodies did not produce detectable signal at the ChP, despite being identified as a top candidate in endothelial cells located at the ChP. This observation suggests that experimental resources are needed to investigate novel genes and proteins of interest such as ANO2.

**Figure 7.**
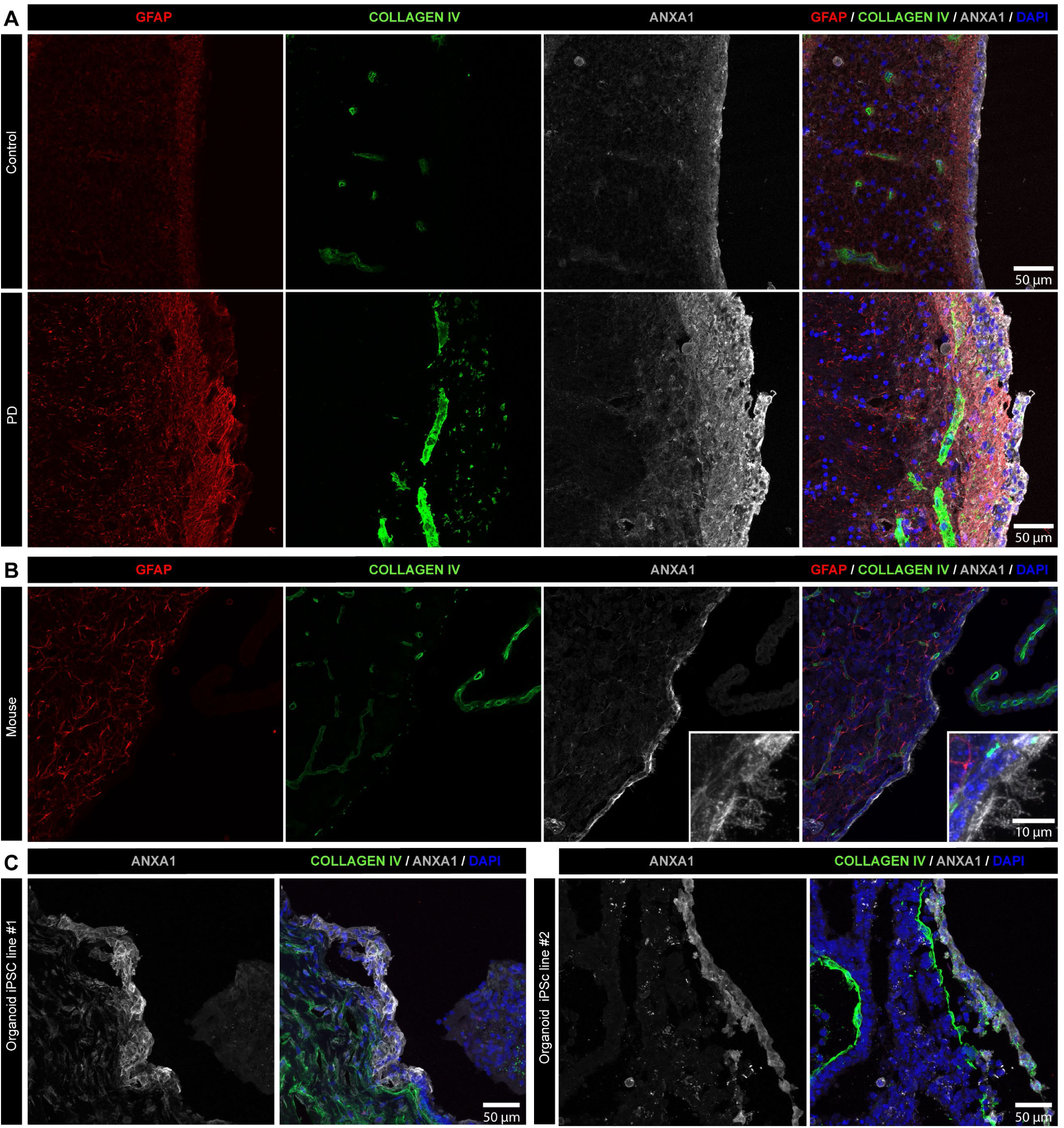
Annexin 1 is localized in ependymal cells located at the border of the ventricle. **A.** Representative confocal images showing GFAP (red), Collagen IV (green) and ANXA1 (white) localized at the ependymal layer forming the border of the lateral ventricle in control and PD postmortem tissues. **B-C.** Confocal images showing GFAP (red), Collagen IV (green) and ANXA1 (white) immunostaining in mouse lateral ventricle (**B**) and ChP organoids generated using two independent iPSC lines (**C**).

**Figure 8.**
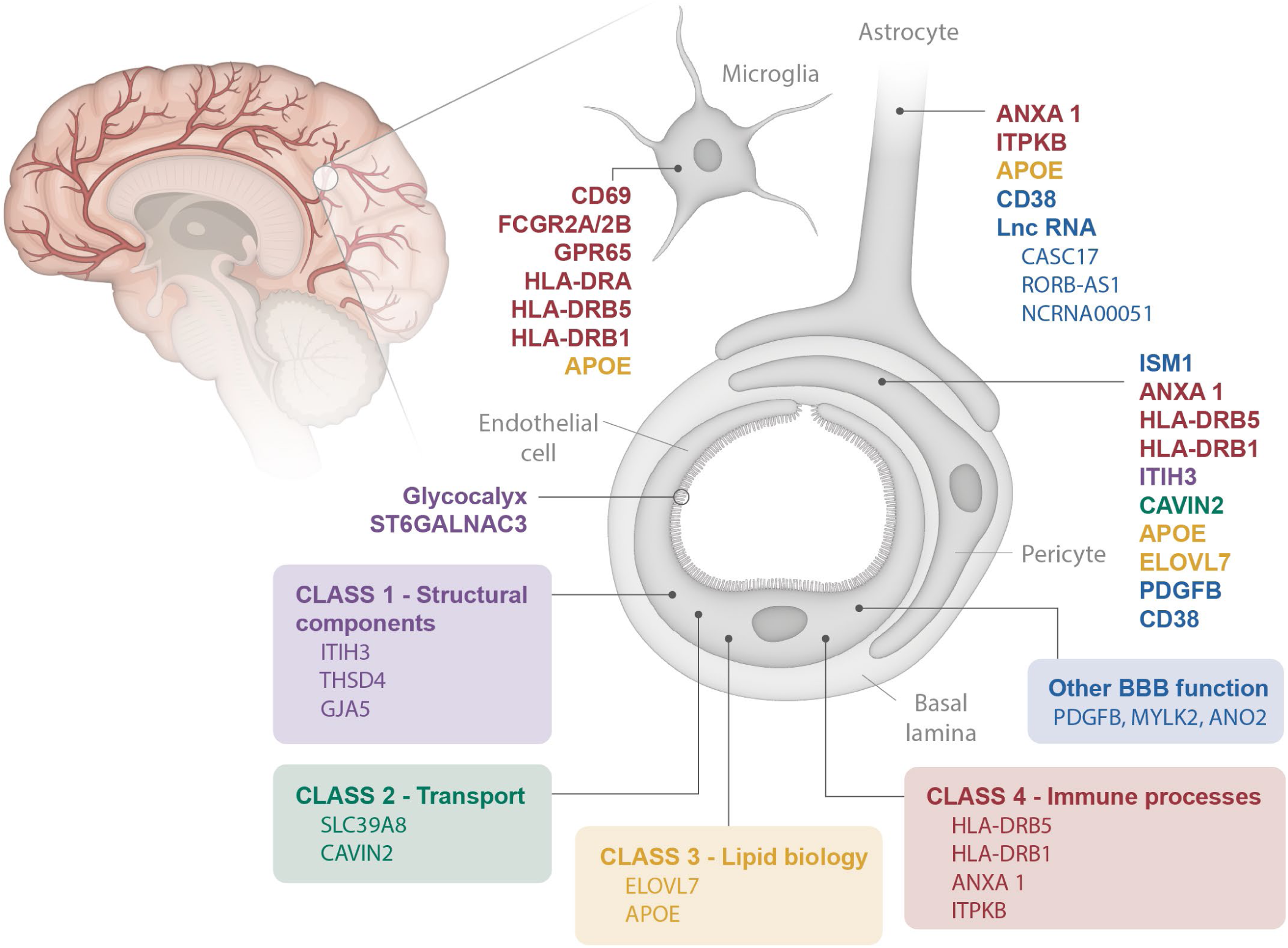
PD risk factors enriched at the neuro-glia-vascular unit by cell type and class of biological function.

## Discussion

This study leveraged published snRNA-seq datasets of human postmortem midbrain, SN, and cortical tissue, followed by an integration with GWAS data to identify PD risk genes associated with cell types that form and maintain functional brain barriers, namely the BBB and BCSFB. The top candidates include genes associated with BBB transcytosis (CAVIN2), BBB protection and anti-inflammation (ANXA1), and the most enriched pathways relate to innate immunity, vascular function, and regulation of cell activation. The investigation was extended to receptor-mediated transcytosis (LRP1), a transport mechanism that regulates molecular exchanges across brain barriers.

### Brain barriers are highly relevant to investigate PD risk factors implicated in protein transcytosis and inflammation

A first key observation resulting from this study is that PD risk factors are expressed by cell types forming brain barriers, including the BCSFB. Previous reports have attempted to associate PD risk loci with specific cell types, and neurons, microglia and oligodendrocytes were the most frequently associated with enriched risk loci and differential expression between PD and control samples^58–60^. For example, a study leveraged mouse single-cell RNA-seq (scRNA-seq) data to propose that dopaminergic neurons, enteric neurons, and oligodendrocytes are associated with PD risk genes, and an analysis of differential gene expression in human PD vs control postmortem brain samples suggested altered GWAS-related gene expression profiles in dopaminergic neurons and oligodendrocytes^61^. In another report, gene enrichment across glial and neuronal cell types of the SN, putamen and frontal cortex included endothelial cells, but did not incorporate pericytes. However, this study compared mouse vs. human transcriptomic datasets and data support the observation of species differences in the enrichment of specific GWAS variants^22^. In another study, cell types of the neuro-glia-vascular unit, including endothelial cells and pericytes, were investigated alongside other glial and neuronal populations but the authors concluded that microglia and oligodendrocytes were the most closely linked cell types to PD^23^. However, a consistent feature of scRNA-seq and snRNA-seq datasets across published studies is the overrepresentation of oligodendrocyte, microglia, and astrocyte nuclei, often accounting for 70-80% of all cells^23^. As a result, it is reasonable to suggest that less abundant cell types, such as endothelial cells and pericytes, may yield lower statistical power in large GWAS-Multi-omic integration studies compared to other populations. Here, we proposed a distinct computational approach in which we calculated cell type enrichment as a ratio of gene expression normalized to neuronal expression. In addition, we focused the calculation of gene enrichment to control samples to avoid potential bias that could be introduced by shifts in cell populations in a disease environment. Furthermore, differential gene expression between control and PD samples was not considered a primary indicator of cell type relevance to enable a scenario where regulation of protein abundance may differ from regulation of gene expression. This strategy revealed candidate genes associated with each cell type of the neuro-glia-vascular unit and the ChP/ependyma. The genes overexpressed in endothelial cells and pericytes related, in part, to innate immunity, vascular function, and regulation of cell activation.

CAVIN2 is a protein involved in the regulation of caveolae biogenesis which, in the context of brain barriers, refers to invaginations at the plasma membrane of endothelial cells that serve as one of the essential regulatory pathways mediating molecular transport at the BBB^62^. Caveolae transcytosis at the BBB is a tightly regulated mechanism, but little is known about the implications of CAVIN2 in these mechanisms. Data provided in this study suggest that patterns of CAVIN2 levels differ between human and mice, as less vascular segments presented with CAVIN2 fluorescent signal in mouse vs. human tissue. However, the human and mouse CAVIN2 are homologous and biological function is therefore likely to be conserved across species. A study proposed that mouse CAVIN2 may regulate caveolae shape, and a *Cavin2* knockout mouse model induced the loss of caveolae formation in the lung and adipose tissue^63^. Interestingly, a study performed in rat brains demonstrated that the endothelial cell efflux pump P-glycoprotein (P-gp) colocalized with CAVIN2 in caveolae-containing fractions, suggesting that rats may be a more appropriate model to study brain CAVIN2, and the protein plays a potential role in P-gp regulatory mechanisms ^64^. Functional P-gp is necessary to prevent the entry of potentially toxic molecules into the brain, and [^11^C]-verapamil-based clinical imaging studies suggested reduced P-pg efflux activity in late-stage people with PD (frontal white matter regions), while P-pg activity appeared increased in patients at early stages (brainstem)^65,66^.

Abundant literature on the role of ANXA1 as an anti-inflammatory and BBB-restorative protein has been published. However, data provided in this study demonstrate the significant discrepancy between mouse and human protein localization in the brain (except at the ependymal layer), and therefore studies that investigated ANXA1 in mouse models may not fully translate to human biology. Nonetheless, these studies support the relevance of the protein in pathophysiological processes relevant to PD and other neurodegenerative diseases. For example, ANXA1 promoted the degradation of amyloid-beta (Aβ) in a murine neuroblastoma cell line^67^, and treatment with human recombinant ANXA1 improved BBB integrity in 5xFAD and Tau-P301L mice, two Alzheimer’s disease models^68^. To the best of our knowledge, the role of ANXA1 at the ependymal layer in health and disease has yet to be explored. In the present study, we did not detect differences in ANXA1 vascular levels between control and PD tissue, but other reports have associated elevated ANXA1 in reactive astrocytes localized at regions of human postmortem brain infarcts^69^. Despite a significant enrichment of ANXA1 at the human brain vasculature, PD-focused studies have mostly investigated the role of ANXA1 in neurons, as several reports identified SOX6+/ANXA1+ dopaminergic neurons selectively vulnerable in PD mouse models^70,71^. In the human SN pars compacta, mRNA-based quantification estimated that approximately 8% of TH+ dopaminergic neurons were also ANXA1+ in control samples, and these populations appeared lost in PD tissue^72^. The MitoPark transgenic mouse model, characterized by the selective depletion of mitochondrial transcription factor A from dopaminergic neurons, recapitulated the selective vulnerability of this cell population^71^, and the manipulation of ANXA1+ dopaminergic neurons induced motor impairments in mouse models^70,71^.

The present study also investigated LRP1 distribution at the control and PD ChP, and confirmed epithelial localization regardless of disease status. This protein has been identified as an α-synuclein receptor^73^, but the role of LRP1 in α-synuclein CSF-to-blood clearance at the ChP remains to be investigated. Published reports suggest that LRP1 may efflux Aβ from the CSF to the blood^74,75^, thereby supporting a similar function towards α-synuclein that could reduce the burden of pathological protein in people with PD. In addition, LRP1 is localized in brain endothelial cells, glia, and neurons, where is regulates α-synuclein spread and transcytosis across the BBB^73,76,77^. The bidirectionality of α-synuclein exchanges at the BBB^78^ along with evidence that LRP1 is implicated in α-synuclein pathology, places brain endothelial cells and ChP epithelial cells in a strong position to serve as potential targets to restore brain homeostasis in PD.

### Gene expression profiles of PD risk factors benefit from protein-level analyses in human postmortem tissues

A second key observation is that gene expression profiles derived from snRNA-seq analyses do not systematically translate to protein abundance in human postmortem brain tissue. This difference highlights potential limitations of single nuclei sequencing, including technical variability and varying sequencing depth that may overlook low abundance transcripts. Validation at the protein level is therefore encouraged to obtain accurate and comprehensive understanding of potential functional outcomes. For example, in this study, we selected CAVIN2 and ANXA1 as two PD risk factors of interest enriched in brain endothelial cells. Transcriptomic data suggested that only a fraction of endothelial nuclei expressed *CAVIN2*, while immunostaining revealed fluorescent signal in most vascular fragments. Similarly to *CAVIN2*, *ANXA1* transcriptomic data poorly reflected protein abundance in human postmortem brain tissue. Gene expression quantifications suggested that ∼20-50% of brain endothelial cells expressed *ANXA1*, while immunostained neurovascular fragments revealed nearly 100% fluorescence signal in the cortex and SN of both control and PD samples. To confirm these observations, immunostaining was performed using an additional antibody for each target. The second CAVIN2 (Proteintech, 12339-1-AP) and ANXA1 (Invitrogen, 71-3400) antibodies provided similar staining patterns. In line with these observations, accumulating studies have been exploring transcript-protein correlations and documented the poor predictive value of transcriptomic studies to estimate protein abundance^79–83^. In the brain, studies observed that correlation may increase significantly for subsets of protein categories (e.g. kinases and membrane-associated proteins^84^, and for specific signaling pathways (e.g. oxidative metabolism and protein synthesis/modification^85^. In a PD-specific context, a study by Zaccaria et al. used laser capture microdissection to extract SN-located dopaminergic neurons from postmortem tissue of control of PD donors, followed by RNA sequencing and nanoliquid chromatography–mass spectrometry to measure transcript and protein abundance, and the authors reported a single transcript-protein pair that were both significantly dysregulated in PD vs control dopaminergic neurons^86^. Furthermore, aging is the highest non-genetic PD risk factor, and a proteomics analyses of the aging cortex identified uncoupled transcript-protein alterations that further complexify the integration of GWAS and transcriptomic datasets to predict the association of cell types to PD pathology. Whether these discrepancies are technical or biological in nature remains to be elucidated^87^, but previously published reports and data presented in this study caution against relying solely on transcriptomic data to conclude on the association of specific cell types with PD risk factors, and suggest that validation of protein expression be included in future studies to offer a comprehensive overview of PD etiology.

### A systematic comparative analysis identified interspecies variation in cell type specificity of selected PD risk factors

A third key observation relates to inter-species differences in protein expression by PD-relevant cell types, which may challenge the translation of research findings documented in mouse models. The most striking observation documented in this study is the abundance of ANXA1 at the human vasculature of the cortex and SN, while the protein is absent at the vasculature of the mouse brain. Instead, and despite 87.5% sequence homology between human and mouse, ANXA1 appeared enriched in dopaminergic neurons of the mouse SN. However, the localization of this protein at the ependymal barrier of the ventricles was conserved across species. The investigation of CAVIN2 revealed that the protein is abundant in human brain endothelial cells, but vascular enrichment was less consistent in mouse brain tissue, and a published study did not detect CAVIN2 in mouse brain by western blot^63^, thus questioning the relevance of mouse models to study these two PD risk factors. In contrast, LRP1 immunostaining confirmed conserved protein expression at both the human and mouse ChP. However, published reports documented lower LRP1 levels in human vs. mouse brain microvessels^88^.Future studies could investigate if LRP1 protein levels are different between mouse and human ChP, as such findings could guide experimental studies aimed at measuring α-synuclein transcytosis or clearance at the BCSFB. The availability of human-based *in vitro* platforms derived from iPSCs provides abundant and renewable material to support investigations of PD risk factors. Here, we confirmed that iPSC-derived astrocytes positively immunostain for ANXA1 and ANO2, and that the pattern of cellular localization is similar to human primary midbrain astrocytes. Furthermore, we generated iPSC-derived ChP organoids and validated LRP1 and ANXA1 protein expression localized at cellular structures reminiscent of their *in vivo* counterparts. Future studies could take advantage of advanced tissue engineering, organ-on-chip, and organoid technologies^89–92^ to propose innovative tools that closely mimic protein expression and distribution observed in the human brain, and increase the translational potential of research studies dedicated to understanding the role of brain barriers in neurodegenerative diseases, including PD.

## Acknowledgements

The authors would like to express their gratitude to the families and people who donated their brains, and therefore enabled the completion of this study. We would also like to express our sincere thanks to Ms. Martine Saint-Pierre for generously sharing her expertise of postmortem immunofluorescence and tissue processing. Funding was provided by the Fondation du CHU de Québec, the Parkinson’s Foundation (Launch Award), and a FRQS Junior 1 Research Scholar Award to AdRJ. OA was supported by a Desjardins and Fondation du CHU de Québec MSc Fellowship. OM was supported by a summer fellowship from the Neuroscience Axis (CRCHUq-UL). MAY was supported by summer fellowships from the Parkinson’s Foundation, MITACS, Sorbonne Université neurosciences. EB is recipient of a Merit award from the Fonds de Recherche en Santé du Québec.

## Authors contributions (using CRediT taxonomy, available at https://casrai.org/credit)

F.B.: Investigation, Methodology, Formal Analysis, Validation, Visualization, Writing—original draft. O.A.: Investigation, Formal Analysis, Visualization, Writing—original draft. V.G.: Formal Analysis, Visualization. L.R.: Investigation, Methodology, Visualization. O.M.: Investigation. M.A.Y.: Investigation. B.J.: Formal Analysis, Methodology, Visualization. L.F.: Methodology. M.B.: Resources. C.B.: Methodology, Supervision. S.L.: Resources, Writing—review & editing. E.B: Writing—review & editing. B.L.: Resources, Methodology, Writing—review & editing. F.C.: Resources, Writing—review & editing. M.P.: Resources, Methodology, Writing—review & editing. A.D.: Supervision, Resources, Writing—review & editing. A.d.R.J.: Conceptualization, Investigation, Formal Analysis, Methodology, Validation, Visualization, Supervision, Funding acquisition, Writing—original draft.

