## Supplementary Material for "Parkinson’s disease risk factors are expressed at brain barriers"

**Supplementary Table 1: List of the marker genes used to separate cell types.**

| NEURO-GLIA-VASCULAR UNIT | CHOROID PLEXUS |
|---|---|
| **Endothelial cells** | **Endothelial cells** |
| *CLDN5*, *PECAM1* | *VWF, ARL15, MECOM, HSP90AA1, ANO2* |
| **Pericytes** | **Epithelial cells** |
| *GRM8*, *PDE7B*, *DLC1*, *SLC6A12*, *TRPC4* | *LINC00276, HRT2C, TMEM72-AS1, GRM8, PCAT1, CP, TTR, SOD3, GPX3, CTSD, BSG, LINGO1, SLC26A3, LINC00486, RAPGEF6, RASGEF1B* |
| **Astrocytes** | **Mesenchymal cells** |
| *SLC1A2*, *GFAP*, *DPP10*, *NRG3*, *AQP4* | *THSD4, SLC4A4, PTGDS, CEMIP, KCNMA1* |
| **Microglia** | **Ependymal cells** |
| *SPP1*, *P2RY12*, *APBB1IP*, *HS3ST4* | *NRXN1, DPP10, CTNNA1, NPAS3, FMN2* |
| **Oligodendrocytes** | **Macrophages** |
| *MBP*, *ST18*, *IL1RAPL1*, *MOBP* | *LRMDA, DOCK4, ARHGAP24, SLC8A1, RUNX1* |
| **Neurons** | |
| *RBFOX3*, *MAP2* | |

**Supplementary Table 2: List of reagents used in this study**

| REAGENT or RESOURCE | SOURCE | IDENTIFIER |
|---|---|---|
| **Cell culture reagents** | | |
| Accutase | Stemcell Technologies | 07922 |
| Accumax | Stemcell Technologies | 07921 |
| Astrocyte Medium | Sciencell | 1801 |
| BSA | BioShop | ALB003.25 |
| FBS | R&D Systems | S11150H |
| TeSR™-E8 | StemCell Technologies | 5990 |
| Geltrex | FisherScientific | A1413302 |
| Matrigel | Corning | 356230 |
| mTeSR Plus medium | Stemcell Technologies | 100-0276 |
| ReLeSR | Stemcell | 100-0483 |
| Triton-X | Sigma | X100-100ML |
| **Antibodies** | | |
| Anti-Annexin 1 | ThermoFisher | 71-3400 |
| Anti-Annexin 1 | Abcam | ab214486 |
| Anti-ANO2 | Bio-techne | NBP2-92888 |
| Anti-ANO2 | Abcam | ab113443 |
| Anti-AQP1 | Santa Cruz Biotechnology | SC-32737 |
| Anti-CD44 | BD Pharmingen | 550538 |
| Anti-Collagen Type IV | MilliporeSigma | AB769 |
| Anti-GFAP | Novus | NBP1-05197 |
| Anti-LRP1 | Abcam | Ab92544 |
| Anti-SDPR (CAVIN2) | Bio-techne | NBP2-57218 |
| Anti- SDPR (CAVIN2) | Proteintech | 12339-1-AP |
| Anti-TH | MilliporeSigma | MAB318 |
| DAPI | Milliporesigma | D9542-1MG |
| Donkey anti-mouse Alexa Fluor 555 | FisherScientific | A31570 |
| Donkey anti-goat Alexa Fluor 488 | FisherScientific | A11055 |
| Donkey anti-rabbit Alexa Fluor 647 | FisherScientific | A31573 |
| Phalloidin-iFluor 488 | abcam | ab176753 |
| **Chemicals, Peptides, and Recombinant Proteins** | | |
| Y-27632 | Selleck Chemicals | S1049 |
| Dextran | Sigma | D8821 |
| Ethanol | Greenfield Global | P025EAAN |
| EGF | ThermoFisher | PHG0311 |
| FGF2 | PeproTech | 100-18C |
| **Experimental Models : Cell Lines** | | |

| | | |
|---|---|---|
| iPSC line #1 | Prof. Dr. Thomas Gasser (Universitätsklinikum Tübingen) and Prof. Dr. Hans R. Schöler (MaxPlanck Institute) | (Reinhardt et al. 2013, *Cell Stem Cell*; PMID: 23472874) |
| iPSC line #2 | Prof. Dr. Tilo Kunath (University of Edinburgh) | (Devine et al. 2011, *Nature Commun.*; PMID: 21863007 |
| **Software and Algorithms** | | |
| GraphPad Prism version 8.0 | https://www.graphpad.com/scientifi c-software/prism/ | |
| Fiji version 2.9.0 | https://imagej.net/software/fiji/downloads | |

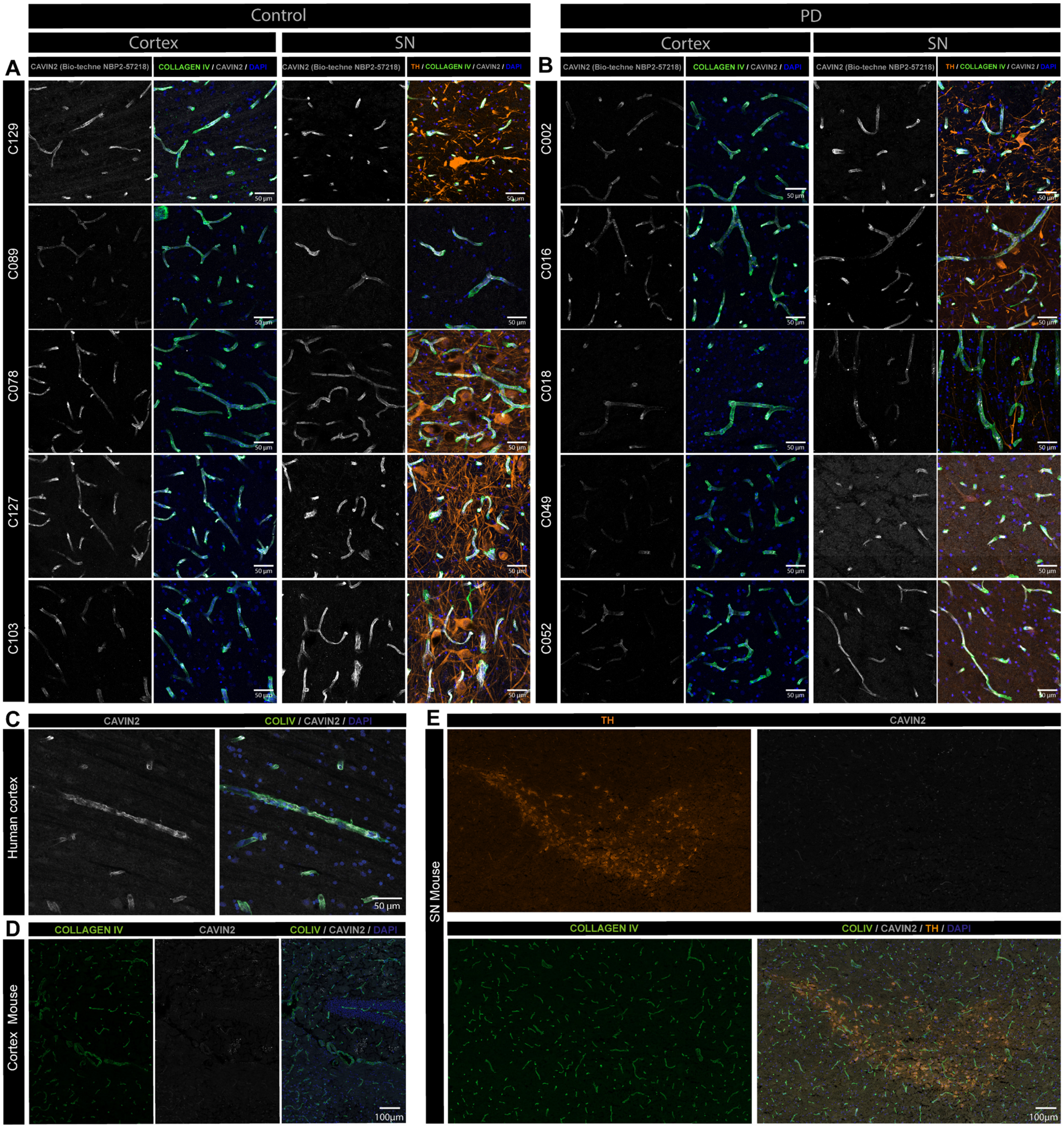


**Supplementary Figure 1. Validation of CAVIN2 signal across different human and mouse postmortem tissues. A-B.** Confocal images showing CAVIN2 (white), Collagen IV (green) and TH (orange) immunostaining in postmortem brain tissues from 5 control (**A**) and 5 PD (**B**)**. C-E.** Confocal images showing CAVIN2 (white), Collagen IV (green) and TH (orange) immunostaining in human postmortem tissue (C) and in mouse cortex (D) and SN (E), using a second antibody for signal validation (Proteintech 12339-1-AP).

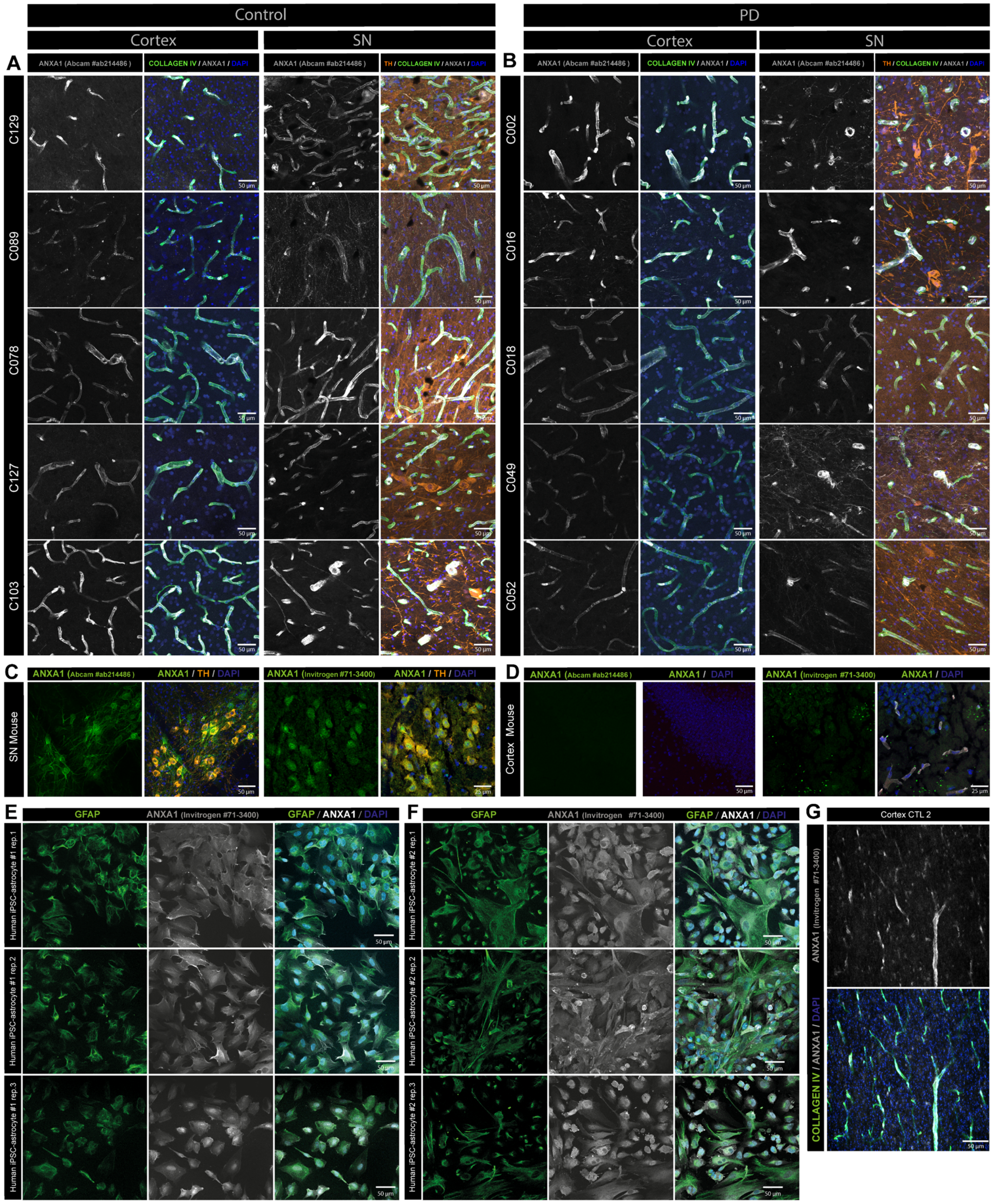


**Supplementary Figure 2**. **Validation of ANXA1 signal across different human and mouse postmortem tissues. A-B.** Confocal images showing ANXA1 (white), Collagen IV (green) and TH (orange) immunostaining in postmortem brain tissues from 5 control (A) and 5 PD (B) donors. **C-D.** Confocal images showing ANXA1 (white), Collagen IV (green) and TH (orange) immunostaining in mouse SN (C) and cortex (D). **(E-F)** Confocal images showing ANXA1 (white) and GFAP (green) immunostaining in iPSC-derived human astrocytes using a second antibody (ThermoFisher/Invitrogen 71-3400). **G.**

Confocal images showing ANXA1 (white) and Collagen IV (green) immunostaining in human cortex using a second antibody to validate signal specificity (ThermoFisher/Invitrogen 71-3400).

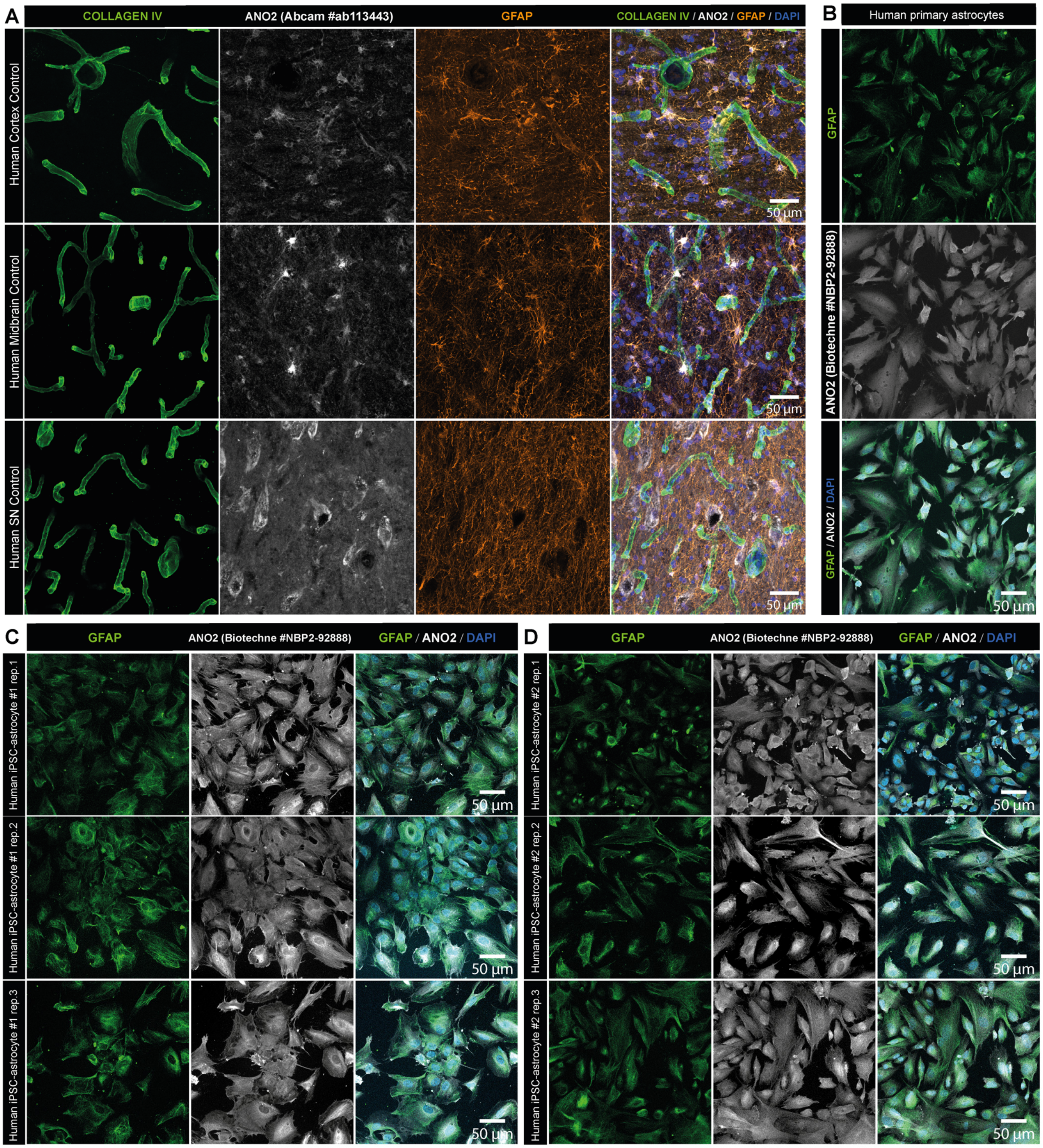


**Supplementary Figure 3. ANO2 is localized in astrocytes in human postmortem cortex. A.** Confocal images showing ANO2 (white), Collagen IV (green) and GFAP (orange) immunostaining in human postmortem cortex from control individuals. In the human postmortem SN, ANO2 localization does not appear vascular nor astrocytic. **B-D.** Confocal images showing ANO2 (white) and GFAP (green) immunostaining in human primary astrocytes (B) and iPSC-derived human astrocytes derived from two independent control donors (C-D). Each panel series in C and D shows immunostaining for three biological replicates produced using iPSC line #1 (C) and #2 (D).

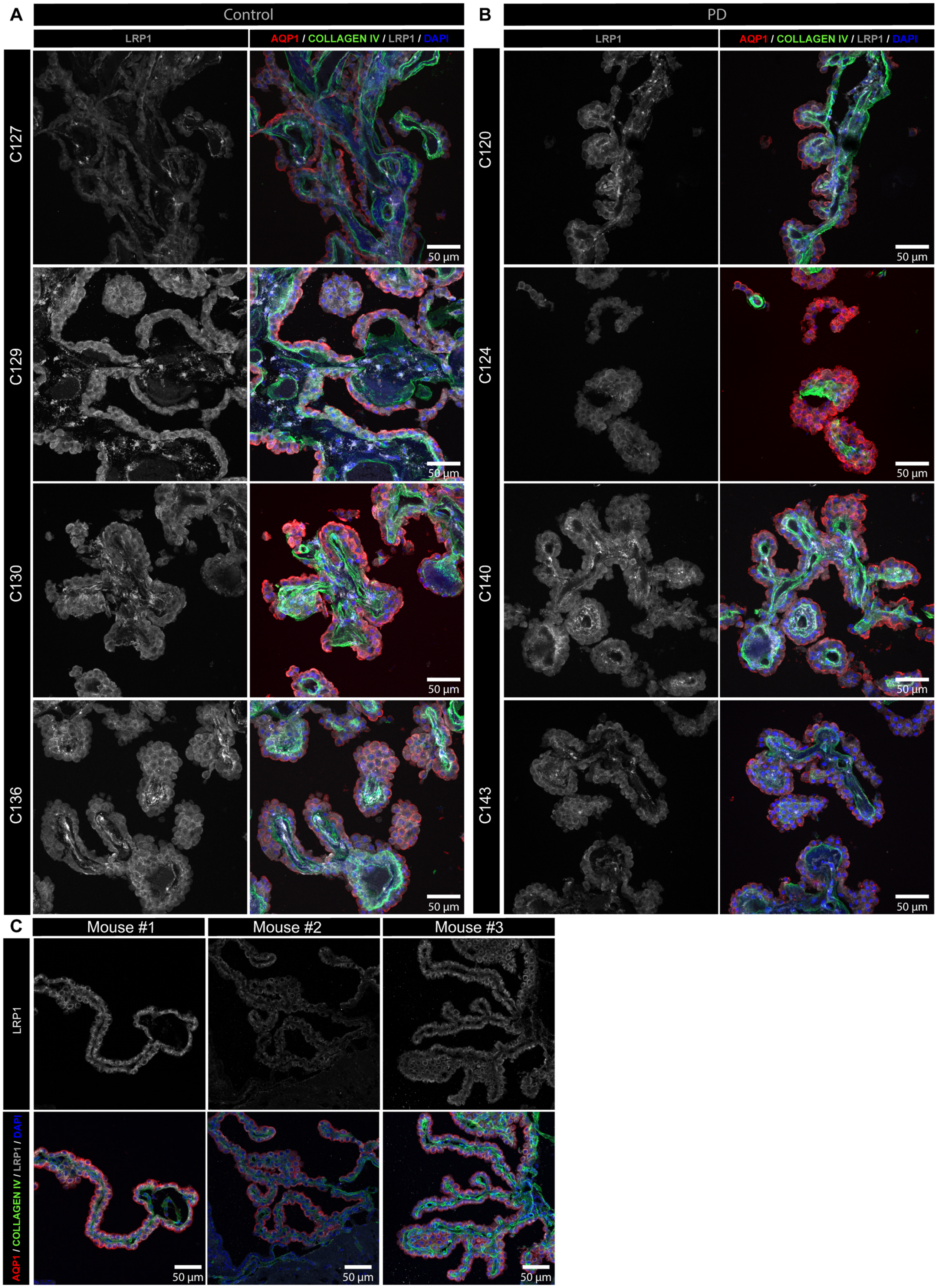
A
Control
LRP1
AQP1 / COLLAGEN IV / LRP1 / DAPI
C127
C129
C130
C136
B
PD
LRP1
AQP1 / COLLAGEN IV / LRP1 / DAPI
C120
C124
C140
C143
50 µm
C
Mouse #1
Mouse #2
Mouse #3
LRP1
AQP1 / COLLAGEN IV / LRP1 / DAPI
50 µm

**Supplementary Figure 4. Validation of LRP1 signal across different human and mouse postmortem tissues. A-C.** Confocal images showing LRP1 (white), Collagen IV (green) and AQP1 (red) immunostaining in ChP from 4 control (A) and 4 PD (B) human postmortem tissues, as well as tissues from 3 different control mice (C).

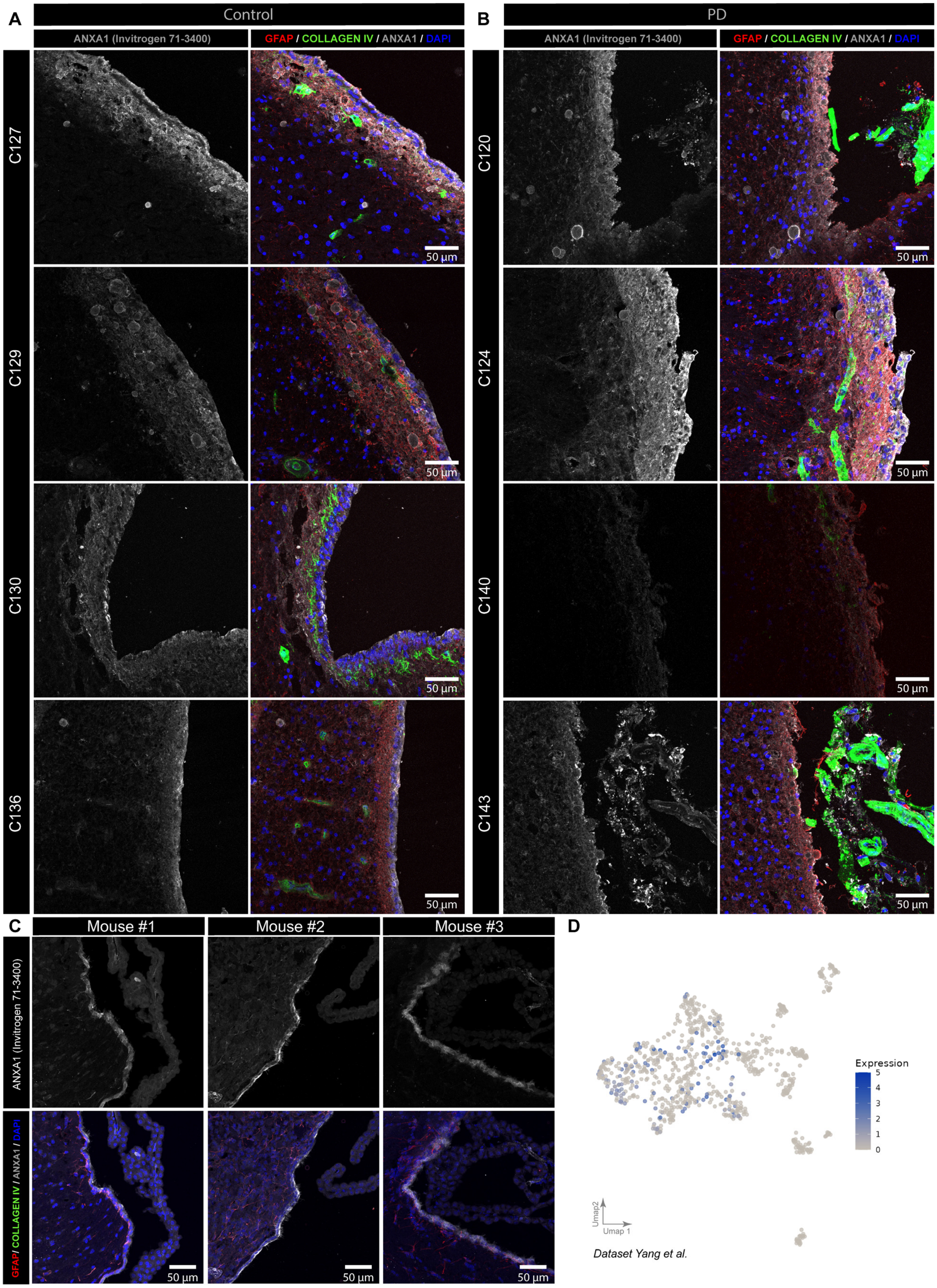

A
Control
ANXA1 (Invitrogen 71-3400)
GFAP / COLLAGEN IV / ANXA1 / DAPI
C127
C129
C130
C136
50 µm
B
PD
ANXA1 (Invitrogen 71-3400)
GFAP / COLLAGEN IV / ANXA1 / DAPI
C120
C124
C140
C143
50 µm
C
Mouse #1
Mouse #2
Mouse #3
ANXA1 (Invitrogen 71-3400)
GFAP/ COLLAGEN IV / ANXA1 / DAPI
50 µm
D
Expression
5
4
3
2
1
0
Umap2
Umap 1
Dataset Yang et al.

**Supplementary Figure 5. Validation of ANXA1 cellular localization in human and mouse ependymal layers. A-C.** Confocal images showing ANXA1 (white), Collagen IV (green) and GFAP (red) immunostaining in ventricles from 4 control (A) and 4 PD (B) human postmortem tissues, as well as tissues from 3 different control mice (C). **D** UMAP represent *ANXA1* expression levels in ependymal nuclei from the Yang et al. dataset.

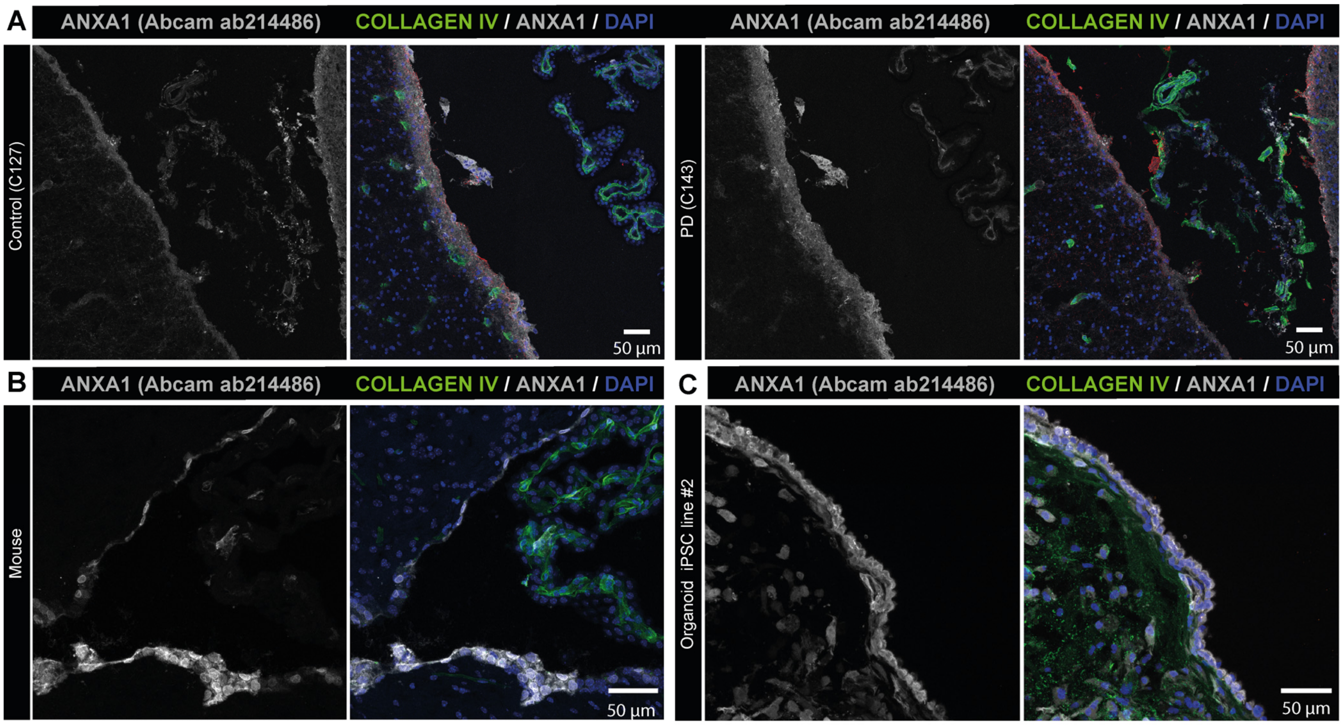


**Supplementary Figure 6. Confirmation of ANXA1 signal specificity using a second antibody. A-C.** Confocal images showing ANXA1(white) and Collagen IV (green) immunostaining using a second ANXA1 antibody (Abcam ab214486) across human postmortem control and PD (A), mouse (B) and ChP organoid samples (C).
